# Phase separation of zonula occludens proteins drives formation of tight junctions

**DOI:** 10.1101/589580

**Authors:** Oliver Beutel, Riccardo Maraspini, Karina Pombo-Garcia, Cécilie Martin-Lemaitre, Alf Honigmann

## Abstract

Tight junctions are cell adhesion complexes that seal tissues and are involved in cell polarity and signalling. Supra-molecular assembly and positioning of tight junctions as continuous networks of adhesion strands is dependent on the two membrane associated scaffolding proteins ZO1 and ZO2. To understand how ZO proteins organize junction assembly, we performed quantitative cell biology and *in vitro* reconstitution experiments. We discovered that ZO proteins self-organize membrane attached compartments via phase separation. We identified the multivalent interactions of the conserved PDZ-SH3-GuK supra-domain as the driver of phase separation. These interactions are regulated by phosphorylation and intra-molecular binding. Formation of condensed ZO protein compartments is sufficient to specifically enrich and localize tight junction proteins including adhesion receptors, cytoskeletal adapters and transcription factors. Our results suggest that an active phase transition of ZO proteins into a condensed membrane bound compartment drives claudin polymerization and coalescence of a continuous tight junction belt.

## Introduction

Tight junctions are cell-cell adhesion complexes that regulate para-cellular flux of solutes and prevent pathogen entry across epithelial and endothelial cell layers including the blood brain barrier (Anderson and Van Itallie, 2009; Citi, 2018). Tight junctions sit at the most apical part of the basolateral plasma membrane and consist of adhesion receptors of the claudin family, which polymerize into a network of intercellular strands creating a selective diffusion barrier (Balda and Matter, 2008; Tsukita et al., 2001). Furthermore, a dense plaque of proteins on the cytoplasmic side regulates junction assembly and provides a connection to the cytoskeleton, polarity proteins, membrane trafficking and transcription (Van Itallie and Anderson, 2014). Decades of genetics, cell biology, and biochemistry have identified the key proteins required for tight junction formation and localization. In addition to adhesion receptors of the claudin family, two homologous scaffolding proteins of the membrane associated guanylate kinases (MAGUK) family (ZO1 and ZO2) are required for tight junction assembly (Fanning and Anderson, 2009; Furuse, 2010; Umeda et al., 2006).

How ZO1 and ZO2 form a membrane attached scaffold that facilitates formation and sub-apical positioning of claudin strands and sequesters cytoskeleton and signalling proteins is not understood on the mechanistic level. As other members of the MAGUK family, ZO proteins contain arrays of conserved protein-protein interaction domains (PDZ, SH3 and GuK) that are connected by mostly unstructured linkers (Funke et al., 2005). This domain organization allows binding to different adhesion receptors including claudins and cytoskeletal proteins simultaneously, which has been suggested to enable crosslinking and scaffolding of junctional proteins with each other and with the actin cytoskeleton (Fanning and Anderson, 2009; Fanning et al., 1998). The scaffolding function of ZOs depend on their ability to oligomerize. ZO proteins form homo and hetero-dimers via their second PDZ domain (Utepbergenov et al., 2006). There is also evidence that additional dimerization can occur independently between SH3 and GuK domains (Umeda et al., 2006), as has been shown for the synaptic ZO homologs (PSD95 and SAP102) (Masuko et al., 1999; Pan et al., 2011). Interestingly, both oligomerization sites are required to form functional tight junction strands (Umeda et al., 2006), which indicates that ZO proteins may form higher oligomers to assemble the junctional plaque. Intriguingly, live cell photo-bleaching experiments revealed that the majority of tight junction plaque proteins including ZO1 and ZO2 are highly dynamic and exchange with the cytoplasmic protein pool within seconds (Garbett and Bretscher, 2014; Shen et al., 2008). Hence, many of the interactions which organize the junctional scaffold are of transient nature. Yet, a mechanism that can explain the supra-molecular assembly of a dense and highly dynamic scaffold that facilitates formation of a continuous tight junction belt is missing.

Recent progress on understanding the nature of non-membrane enclosed compartments has revealed that many of these highly dynamic assemblies form via liquid-liquid phase separation of scaffolding proteins (Brangwynne et al., 2009; Li et al., 2012). Phase separation of proteins into condensed liquid states is facilitated by weak multivalent protein-protein or protein-RNA interactions (Pak et al., 2016; Wang et al., 2018). Prominent examples are P granules, nucleoli or stress granules [17]. Importantly, phase separation has also been found to underlie the assembly of adhesion complexes related to tight junctions such as the immunological synapse and the post-synaptic density (Su et al., 2016; Zeng et al., 2016). Interestingly, ZO proteins, the scaffolders of the tight junction, are homologs of the postsynaptic MAGUK protein PSD95, which organizes the post-synaptic density via phase separation. As PSD95, ZO proteins contain multivalent protein-protein interaction domains. In addition, ZOs also contain unique and long intrinsically disordered domains. These structural features of ZO proteins together with the known properties of the tight junction plaque made us hypothesize that phase separation might play a role in the formation of tight junctions.

Here, we present evidence, based on quantitative cell biology and *in vitro* reconstitution, that ZO proteins form condensed liquid-like compartments *in cellulo* and *in vitro* via phase separation. We further identified the domains that drive and regulate phase separation and show specific partitioning of tight junction proteins into the condensed scaffold. Our results suggest that phase separation of ZO proteins into a condensed membrane bound phase underlies partitioning and possibly nucleation of claudin and actin polymerization and hence tight junction formation.

## Results

### Quantification of ZO1 and ZO2 concentrations and dynamics in MDCK-II cells

The scaffolding proteins ZO1 and ZO2 are necessary to assemble functional tight junctions in epithelial cells (Umeda et al., 2006). It has been shown that ZO1, while being highly enriched at the tight junction, rapidly turns over with the cytoplasmic pool (Shen et al., 2008; Yu et al., 2010) (Figure 1A). We used these studies as our starting point, to ask whether the concentrations and dynamics of endogenous ZO1 and ZO2 at epithelial tight junctions support the hypothesis of phase separation of ZO1 and ZO2.

**Figure 1.**
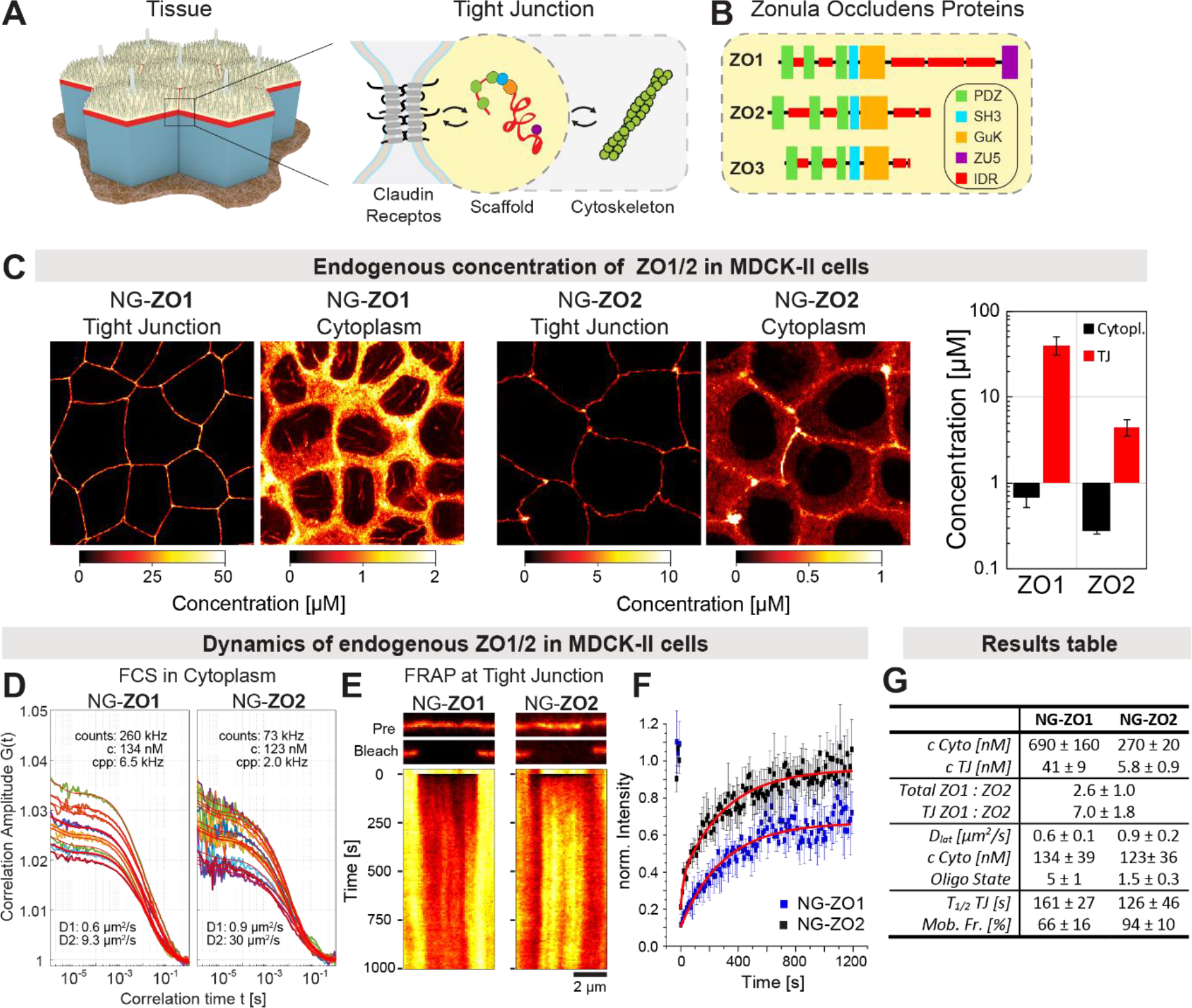
Quantification of ZO1 and ZO2 concentrations and dynamics in MDCK-II cells. (A) Scheme of the tight junction complex (TJ) in mammalian epithelial cells. The TJ complex consists of adhesion receptors, scaffolding proteins and cytoskeletal elements. (B) The domain structure of the main TJ scaffolding proteins (ZOs). All three ZO homologs belong to the MAGUK family and contain multiple protein-protein interaction domains and extended intrinsically disordered linkers. (C) Quantification of endogenous concentration of ZO1 and ZO2 in the cytoplasm and the TJ of MDCK-II cells. mNeonGreen (NG) was inserted N-terminally using CRISPR (S1A) and quantitative fluorescence microscopy was used to determine concentrations from confocal images. Calibration of NG fluorescence intensity is shown in S1C. Both ZO1/2 are strongly enriched at the TJ and are present at sub-micromolar concentration in the cytoplasm. ZO1 concentration is 2.6-times higher in the cytoplasm and 7-times higher at the TJ compared to ZO2. The results are summarized in table (G) (n = 30 cells, ± SD). (D) Quantification of dynamics of NG-ZO1/2 in the cytoplasm using FCS. Shown are FCS-fits from 10 different cells for NG-ZO1 and NG-ZO2. The results show that both NG-ZO1/2 have a slow diffusing fraction, which indicates binding to larger complexes in the cytoplasm. Based on the comparison of the single particle brightness (cpp) of free NG and NG-ZO1/2, the oligomeric state of NG-ZO1 and NG-ZO2 was 5 and 1.5, respectively. The results are summarized in (G). (E) Quantification of ZO1/2 dynamics at the TJ using FRAP. NG-ZO1/2 was bleached selectively at the TJ and recovery was measured at room temperature over time. Kymographs show that ZO1/2 recovered rapidly from the cytoplasm but not from the adjacent junctional regions. (F) Double exponential fit of normalized and averaged recovery curves of 5 independent measurements (± SD). (G) Summary of quantitative imaging, FCS and FRAP.

We used CRISPR/Cas9 to generate homozygous N-terminal insertions of the fluorescence protein mNeonGreen (NG) in the ZO1 and ZO2 *loci* in MDCK-II cells, respectively (Figure S1A,B). Next, we used quantitative microscopy to determine the endogenous cytoplasmic and junctional concentration of NG-ZO1 and NG-ZO2 (Figure 1B). The quantification was based on calibrating the image intensities with purified NG (Figure S1C). We found an average cytoplasm concentration of c_ZO1-Cyto_ = 0.8 μM, c_ZO2-Cyto_ = 0.3 μM and we estimated the junctional concentration to be c_ZO1-TJ_ = 30 μM, c_ZO2-TJ_ = 4 μM. The quantification allowed us to calculate the ratio of ZO1:ZO2 in the cytoplasm and at the junction. We found that ZO1 was 2.5-fold more abundant in the cytoplasm. Interestingly, the ratio increased to 7-fold at the tight junction, indicating a preferential binding of ZO1 over ZO2 to the junction. To reinforce the imaging results, we used quantitative PCR to determine the average amount of mRNA transcripts of ZO1, ZO2 and ZO3 in wild type MDCK-II cells. In line with the imaging results, qPCR showed that the total ZO2 mRNA levels was 3-fold reduced compared to ZO1. mRNA of ZO3, the third ZO homolog, which is not required for tight junction formation, was only expressed at 2% compared to the level of ZO1 (Figure S1D).

Next, we used fluorescence correlation spectroscopy (FCS) and fluorescence recovery after photobleaching (FRAP) to measure and compare the dynamics of endogenous NG-ZO1 and NG-ZO2 in the cytoplasm and at the tight junction. FCS measurements in the cytoplasm provided an independent estimate of the average NG-ZO1 and NG-ZO2 concentration, which were consistent with but slightly lower than the quantitative imaging results (Figure 1D, G). FCS analysis in the cytoplasm identified two diffusive components for both ZO1 and ZO2 with lateral diffusion coefficients of D1_ZO1_ = 0.6 μm^2^/s, D2_ZO1_ = 9.3 μm^2^/s and D1_ZO2_= 0.9 μm^2^/s, D2_ZO2_ = 30 μm^2^/s. The fast diffusion fraction, which accounted only for 20% of the total fraction, can be attributed to NG-ZO1 and NG-ZO2 diffusing as monomers in the cytoplasm. The slower component indicates that 80% of ZO1 and ZO2 moved as larger complexes in the cytoplasm. Interestingly, comparing the molecular brightness of free NG in MDCK-II cells to NG-ZO1 and NG-ZO2, we found that NG-ZO1 and NG-ZO2 brightness were 5-fold and 1.5-fold higher than monomeric NG, respectively. This indicates that ZO1 and ZO2 form oligomers in the cytoplasm with distinct stoichiometry. To measure the dynamics of NG-ZO1 and NG-ZO2 at the tight junction, we switched to FRAP since FCS is limited to measuring relatively fast dynamics. We bleached regions selectively at the tight junction and measured the fluorescence recovery over time. In line with previous reports on FRAP of heterologously expressed ZO1 (Shen et al., 2008; Yu et al., 2010), we found that endogenous NG-ZO1 recovered to 70% of its initial concentration at 23°C with a time constant of t_1/2_ = 161s. The recovery came predominantly from the cytoplasm and not from the adjacent junctional area. Interestingly, NG-ZO2 recovery was slightly faster with t_1/2_ = 126s and an immobile fraction was absent. This shows that ZO2 interactions at the tight junction are more transient than ZO1.

Our data on the quantification of the concentration and dynamics of endogenous ZO1 and ZO2 so far are in line with previous studies that used heterologously expressed ZO1 (Shen et al., 2008; Yu et al., 2010). Studying endogenous ZO1/2 we found that in MDCK-II cells ZO1/2 are present at sub-micromolar levels in the cytoplasm, are enriched up to 80-fold at the tight junction and the junctional pool exchanges with the cytoplasmic pool within seconds to minutes. ZO1 is the dominant ZO homolog at the tight junction and its dynamics are reduced compared to ZO2. These features, in particular the highly condensed state at the tight junction and its dynamic turn over with a lower cytoplasmic pool, are signatures of liquid phase separation of proteins (Banani et al., 2017; Berry et al., 2018). This in addition to the known multi-domain structure of ZO proteins with its extended intrinsically disordered linker regions (Harmon et al., 2017), further motivated us to test the hypothesis that the ZO1/2 junction scaffold may assemble via a phase separation process.

### ZO1 and ZO2 phase separate into liquid membrane attached compartments in cells

To test our hypothesis that ZO proteins may form phase separated compartments we transiently expressed all ZO homologs (ZO1, ZO2 and ZO3) with a N-terminal Dendra2 tag in MDCK-II, respectively (Figure 2A). At low expression levels all constructs localized to the tight junction belt (Figure 2A upper panel). However, we noticed that in cells expressing ZO proteins at higher concentrations bright non-junctional assemblies were visible (Figure 2A middle panel). Time-lapse imaging revealed fusion and fission of these assemblies within seconds, which indicated liquid-like material properties (Figure 2A lower panel). While ZO3 formed perfectly spherical droplets in the cytoplasm, ZO1 and ZO2 assemblies were attached to the cell membrane and often fused into continuous domains at the membrane interface. Overexpression of other multi-domain scaffolding proteins (MPP5, DLG1, MAGI3, MPDZ) N-terminally tagged with Dendra2 or Dendra2 alone showed homogenous distributions (Figure 2B, S2A,B), indicating that formation of assemblies is rather specific to ZO proteins. Exchanging the fluorescent tag on ZO2 to CLIP-tag resulted in similar assemblies as in Figure 2A, indicating no significant influence of the tag on protein assemblies (Figure S2B). While the formation of large-scale ZO assemblies was obviously induced by the overexpression, we speculated that these experiments revealed an intrinsic capacity of ZO proteins to phase separate into liquid-like membrane attached compartments and that this could be an important function to facilitate junction formation.

**Figure 2.**
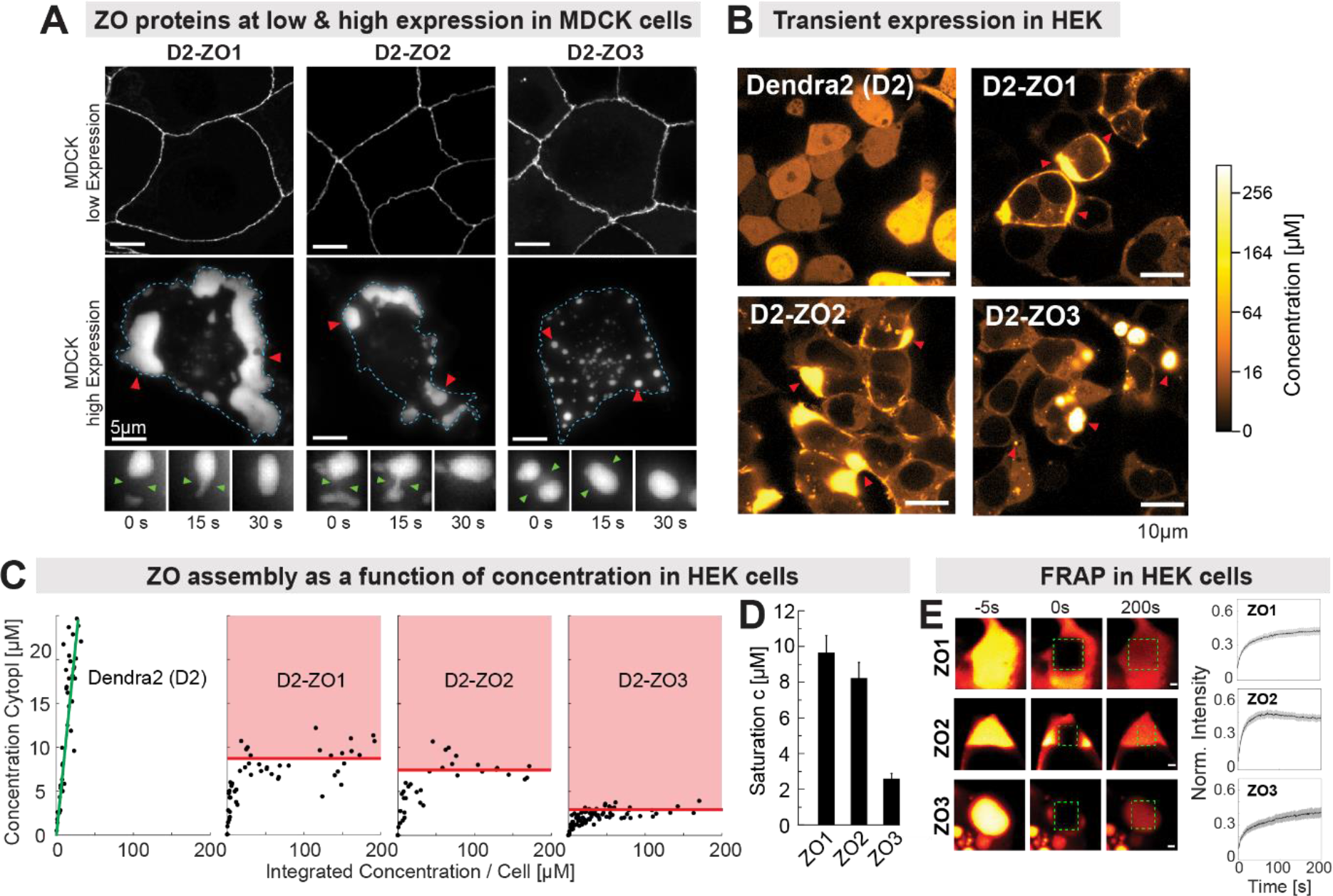
ZO proteins phase separate into liquid membrane attached compartments in cells. (A) Transient expression of N-terminal Dendra2 (D2)-tagged ZO1/2/3 in MDCK-II cells. At low expression levels all constructs were enriched at the TJ (upper panel). At higher expression bright assemblies appeared which were attached to the cell membrane for ZO1/2 or cytoplasmic in case of ZO3 (red arrows, middle panel). ZO assemblies fused into larger structures over time, indicating liquid-like properties (lower panel). (B) Transient expression of D2-ZO1/2/3 in HEK293 cells confirmed formation of assemblies seen in MDCK-II cells (red arrows). To determine at which concentration ZO assemblies form, we used quantitative microscopy to measure the cytoplasmic and total concentration of individual HEK293 cells expressing the constructs at random concentrations. To visualize the low cytoplasmic concentrations, the intensity scale is non-linear (gamma = 0.5). (C) Quantification of cytoplasmic and total concentration of over 50 individual HEK293 cells expressing D2-ZO1/2/3, respectively. Plotting the cytosolic concentration against the total concentration of HEK293 cells expressing Dendra2 alone shows a linear relation (green line), i.e. the more protein is expressed the higher is its cytoplasmic concentration. In contrast, for HEK293 cells expressing D2-ZO1/2/3, a clear saturation of cytoplasmic concentration is seen. Above the saturation level the additional protein is segregating into condensates and the cytoplasmic concentration is hardly changed. This behaviour is characteristic for a phase transition. (D) Quantification of the saturation concentration from the graphs in (C). D2-ZO1/2 have comparable saturation concentrations around 9-8 μM, while D2-ZO3 has a much lower saturation concentration of 2 μM (± SD). (E) FRAP measurements of D2-ZO1/2/3 in the condensed phase in HEK293 cells. All constructs showed rapid recovery, indicating diffusion inside the droplets and exchange with the cytoplasmic pool. Recovery curves are mean of 5 droplets (shadow shows SD).

To determine the concentration at which ZO condensates spontaneously assemble, we determined the ZO phase separation as a function of protein concentration in cells. To this end, we chose HEK293 cells since they do not make tight junctions and have a high transfection efficiency. ZO protein constructs were expressed in a wide range of concentrations due to the stochastic nature of the plasmid transfection process. Similarly to MDCK-II cells, bright liquid-like condensates were clearly visible in HEK293 cells (Figure 2B). We then used quantitative fluorescent microscopy to determine the cytoplasmic concentration of ZO proteins as a function of the total expression level per cell (Figure 2B). Control proteins, including other MAGUK and PDZ-scaffolding proteins (MPP5, DLG1, MAGI3 MPDZ and Dendra2) did not form condensates and displayed a linear relation between cytoplasmic and total concentration over a large concentration range (Figure 2C, S2A-C). In contrast, the cytoplasmic concentration of ZO homologs rapidly saturated when expression levels reached the low micro-molar regime. At expression levels exceeding the cytoplasmic saturation concentration, bright condensates appeared and grew in size with increasing expression levels. We then determined the saturation concentrations of the phase transition by linear fitting of the plateaus in Figure 2C. We found that ZO1 and ZO2 had comparable saturation concentrations with c_sat-ZO1_ = 9.5 μM and c_sat-ZO2_ = 8 μM, while ZO3 phase separated at a significantly lower concentration c_sat-ZO3_ = 2.1 μM (Figure 2D).

To determine the dynamics inside ZO condensates and compare them to ZO dynamics at the tight junction we used FRAP (Figure 2E). We partially and fully bleached ZO condensates and measured recovery times (Figure 2G). All ZO condensates showed rapid recovery within seconds indicating liquid-like material properties.

Taken together, the overexpression experiments indicate that ZO proteins have an intrinsic capacity to form condensed compartments via liquid-liquid phase separation in epithelial cells. Comparison of endogenous concentrations to the concentrations required for spontaneous phase separation showed that the endogenous cytoplasmic concentrations of ZO1/2 are below the phase separation threshold. However, at the tight junction the ZO1/2 concentration is above the threshold. Therefore, ZO1/2 are expected to form a condensed phase at the tight junction. Comparison of ZO dynamics determined by FRAP at the tight junction (Figure1E, F) and in phase-separated droplets (Figure 2E) showed that ZO1/2 dynamics at the tight junction are significantly slower and constrained than predicted for a liquid phase with the properties determined in phase-separated ZO1/2 droplets in HEK293 cells. This indicates that additional interactions take place at the mature tight junction, which are not present in the condensed ZO droplets. For example, lateral diffusion of ZO1/2 may be restricted by the underlying network of claudin polymers and other tight junction scaffolding proteins (MUPP1, PAR3) and F-actin may further cross-link the ZO1/2 scaffold (Shen et al., 2008; Yu et al., 2010).

### Purified ZO proteins form phosphorylation sensitive liquid condensates *in vitro*

Based on our previous experiments we speculated that ZO proteins have an intrinsic capacity to form liquid-like condensates in cells. To test whether condensation is indeed directly mediated by ZO proteins or if other cellular players are involved we set up a series of *in vitro* experiments. We used a eukaryotic insect cell system to express and purify full-length ZO1, ZO2 and ZO3 with C-terminal mEGFP and MBP-tag (Figure 3A). After cleavage of the MBP tag, the purified ZO proteins were soluble up to high micro-molar concentrations in high salt buffers (500 mM KCl, pH 8). However, when we changed the buffer to physiological salt concentration (150 mM KCl, 2% PEG-8k, pH 7.2) we observed spontaneous condensation of ZO1, ZO2 and ZO3 into liquid droplets that fused over time, showing that these proteins are sufficient to form condensates via liquid-liquid phase separation (Figure 3B, C). Using this condition, we determined the affinity for ZO1, ZO2 and ZO3 to phase separate *in vitro* (Figure 3D). In line with the *in vivo* observations, ZO3 had the strongest affinity to phase separate (c_sat_ = 1.2 μM), ZO2 the second strongest (c_sat_ = 3 μM) and ZO1 the lowest (c_sat_ = 4.3 μM).

**Figure 3.**
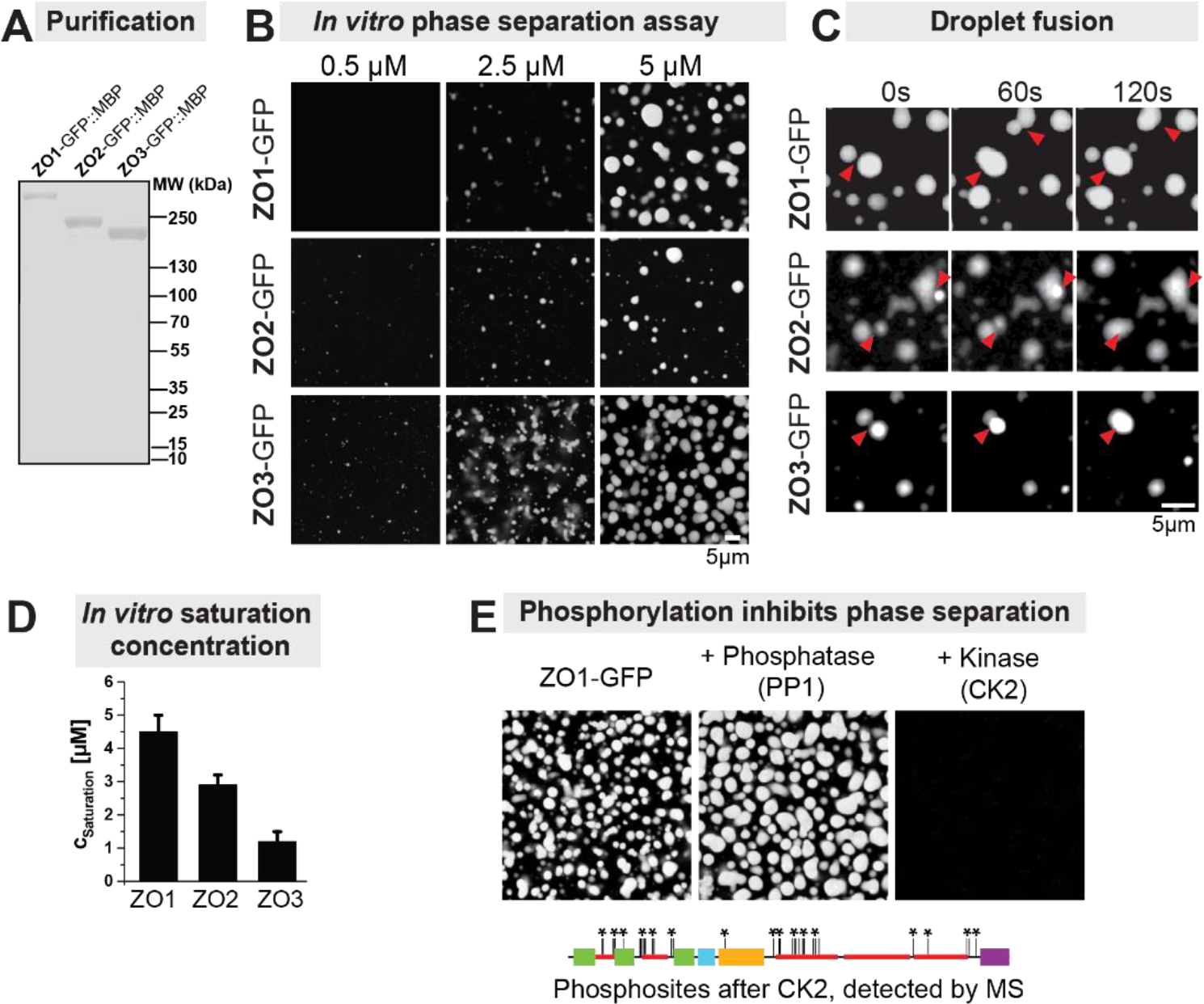
Purified ZO proteins form phosphorylation sensitive liquid condensates *in vitro*. (A) SDS-PAGE showing purified ZO1/2/3 – GFP before removal of the MBP-tags. (B) Concentration-dependent phase separation into liquid droplets of ZO1, ZO2 and ZO3 *in vitro* (Buffer: 150mM NaCl, 20mM TRIS, pH 7.4, 3% PEG(8k)) (C) Fusion events of droplets over time indicate liquid-like material properties of ZO condensates. (D) Saturation concentrations, i.e. concentration outside of the condensed phase, *in vitro*. The same trend was observed as *in vivo*: ZO3 had the highest affinity to phase separate, ZO1 the lowest (n = 3 experiments, ± SD). (E) De-phosphorylation by PP1 promoted phase separation. Phosphorylation of ZO1 by CKII inhibited phase separation. We detected 38 phosphosites (asterisk) after phosphorylation ZO1 by CKII using mass spectrometry.

ZO1/2 are known to be phosphoproteins and their phosphorylation state has been previously linked to the tight junction formation (Dörfel and Huber, 2012). Enhanced phosphorylation of tight junction proteins including ZO1/2 reduces tight junction formation (Rao et al., 2002; Sallee and Burridge, 2009). To test whether phosphorylation state affects the phase separation ability of ZO1, we phosphorylated and dephosphorylated the protein *in vitro* using casein kinase-2 (CK2) and the phosphatase PP1. We found that dephosphorylated ZO1 efficiently phase separated into liquid droplets. In contrast, phosphorylated ZO1 was unable to phase separate under the tested conditions (Figure 3E). Mass spectrometry revealed that CK2 phosphorylated ZO1 at 38 residues. While there seemed to be a phosphorylation hotspot in the disordered U6 region downstream of the GuK domain, a dedicated study is required to understand if phosphorylation of specific residues is sufficient for inhibiting phase separation.

Taken together, the *in vitro* results confirmed that ZO proteins have an intrinsic capacity to phase separate into liquid compartments and we observed similar difference in affinities for phase separation between the homologs (ZO3 > ZO2 > ZO1) as seen in HEK293 cells. The phase transition of ZO1 can be modulated via its phosphorylation state. Hence, de-/phosphorylation could be a mechanism to actively trigger phase separation of ZO1/2 in a locally controlled manner.

### Multivalent *cis* and *trans* protein-protein interactions drive phase separation of ZO proteins

Liquid-liquid phase separation of proteins is known to be driven by transient, multivalent protein-protein interactions (Li et al., 2012; Pak et al., 2016; Wang et al., 2018). ZO proteins contain a number of conserved protein-protein interaction domains which are connected by unique intrinsically disordered domains (U1-6) (Figure 4A). Based on previous biochemical studies on ZO1 and other MAGUK proteins, we speculated that three regions of ZO proteins contribute valences required for phase separation. The second PDZ domain (PDZ2) at the N-terminus induces homo- and hetero-dimerization with the other ZO homologs (Utepbergenov et al., 2006; Wu et al., 2007). The conserved core of the ZO proteins, the PDZ3-SH3-GuK supra-domain, has been proposed to oligomerize potentially via domain swapping (Umeda et al., 2006; Ye et al., 2018). Two regions, the U6 domain and the far end of the intrinsically disordered C-terminal regions of ZO1 and ZO2, have been proposed to bind back to the PSG supra-domain (Fanning et al., 2007; Lye et al., 2010; Spadaro et al., 2017). Additionally, intrinsically disordered regions of ZOs may drive phase separation via low affinity interactions (Wang et al., 2018).

**Figure 4.**
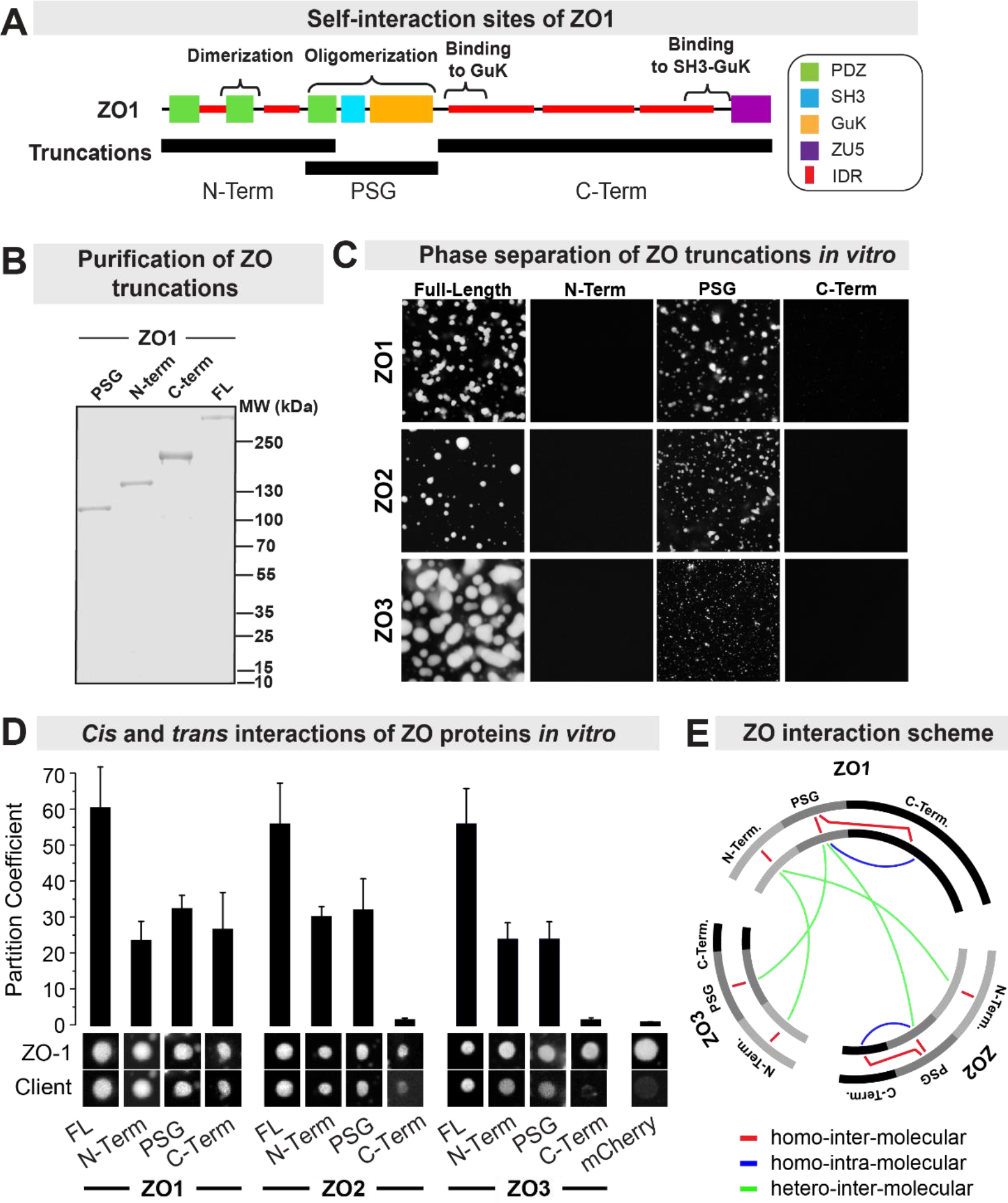
Multivalent protein-protein interactions drive phase separation of ZO proteins. (A) Scheme of known self-interaction sites of ZO1 and the truncations we tested. (B) To find the domains responsible for phase separation *in vitro,* we express and purified three truncations (N-term, PSG and C-term) for ZO1, ZO2 and ZO3. (C) Phase separation assay of truncation mutants of ZO1, ZO2 and ZO3 *in vitro* (same buffer as in 3B, protein concentrations 5μM). For all ZO homologs the full-length protein and the PSG fragment consistently phase separated into liquid like droplets under the tested conditions. The N-Term and C-Term fragments did not phase separate. (D) Partitioning assay to determine protein-protein interactions of phase separated full-length ZO1 with fragments of ZO1, ZO2 and ZO3 (Clients). Partitioning was determined by computing the ratio of fluorescence inside to outside of the droplet for the client proteins. (n > 10 droplets, ± SD). Partitioning of ZO2 and ZO3 is shown in figure S4A. (E) Interaction scheme of ZO proteins based on the partitioning assay. Linear sequence of ZO1, ZO2 and ZO3 and its N-Term, PSG, C-Term regions are depicted as two circles to indicate protein-protein interactions between the same homolog (red), between different homologs (green) and for intra-molecular interactions (blue).

To identify the domains required for phase separation, we expressed three fragments of the ZO proteins and tested them using the *in vitro* phase separation assay (2% PEG-8k, 150 mM KCl, pH 7.2, Figure 4B, C). We found that the purified N-terminal fragments of ZO1/2/3 containing the PDZ1, PDZ2 and PDZ3 domains did not phase separate under these conditions. The C-terminal mostly disordered fragment did not phase separate either. But the conserved core of MAGUK proteins containing the PDZ3-SH3-GuK (PSG) supra-domain showed condensation into liquid droplets. This result indicates that the PSG supra-domain is essential for condensation of ZO proteins into liquid compartments. However, the saturation concentration of the PSG fragment was higher compared to the full-length ZO1, which indicates that additional interactions outside the PSG domain are important.

To test for interactions of ZO proteins with itself (*cis*) and between the other homologs (*trans*) we set up a two-colour partitioning assay *in vitro* (Figure 4D, S4A). We used the respective full-length (FL) ZO protein labelled with mEGFP to form condensed droplets and then measured the partitioning of mCherry-labelled ZO homologs and its N-terminal, PSG and C-terminal fragments into the condensed phase. The partitioning coefficient directly reports on positive (enrichment) or negative (exclusion) interactions and therefore allows us to map *cis* and *trans* interactions of ZO proteins. The results revealed that both the N-terminal and the C-terminal fragments of ZO1 and ZO2 bind to their respective full-length homologs, which confirms previous biochemical protein interaction studies (Fanning et al., 1998; Spadaro et al., 2017). Interestingly, while the N-terminus of ZO1 can interact in *trans* with ZO2 or ZO3, the interactions of the C-termini are restricted to *cis* interactions, e.g. ZO1-C-term can bind only to ZO1-FL and similarly ZO2-C-term can only bind to ZO2-FL. Another surprising result was that ZO2 and ZO3 do not interact with each other in our partitioning assay (Figure S4A). We summarized the multivalent *cis* and *trans* interactions of the ZO homologs in Figure 4E, which also highlights the central role of ZO1 in the ZO interactome. Taken together, the results of the ZO self-interaction measurements support the idea that phase separation of ZO proteins is driven by transient multivalent interactions between the same homolog (*cis*) and between different homologs (*trans*).

### The condensed phase of ZO proteins selectively sequesters tight junction proteins

ZO1/2 have been proposed to recruit and organize other proteins at the tight junction (Figure 5A). However, interactions of ZO proteins with other junctional proteins (clients) were reported to be of relatively low affinity. For instance, the PDZ domains of ZO1 interact with the C-terminal tails of adhesion receptors with affinities ranging between K_d_ = 1 - 50 μM (Itoh et al., 1999; Nomme et al., 2015; Pereda et al., 2008). It is therefore unclear how stable interactions between the scaffolding protein and its clients are achieved at endogenous expression levels. Phase separation may provide a solution to this problem. Because phase separation produces compartments with a high internal concentration of ZO proteins, low affinity interactions (with clients) will result in efficient binding within the phase but not outside. This mechanism should result in a strong up-concentration (partitioning) of clients into the phase. Hence, ZO phase separation may provide a way to selectively enrich junctional components in a dedicated compartment.

**Figure 5.**
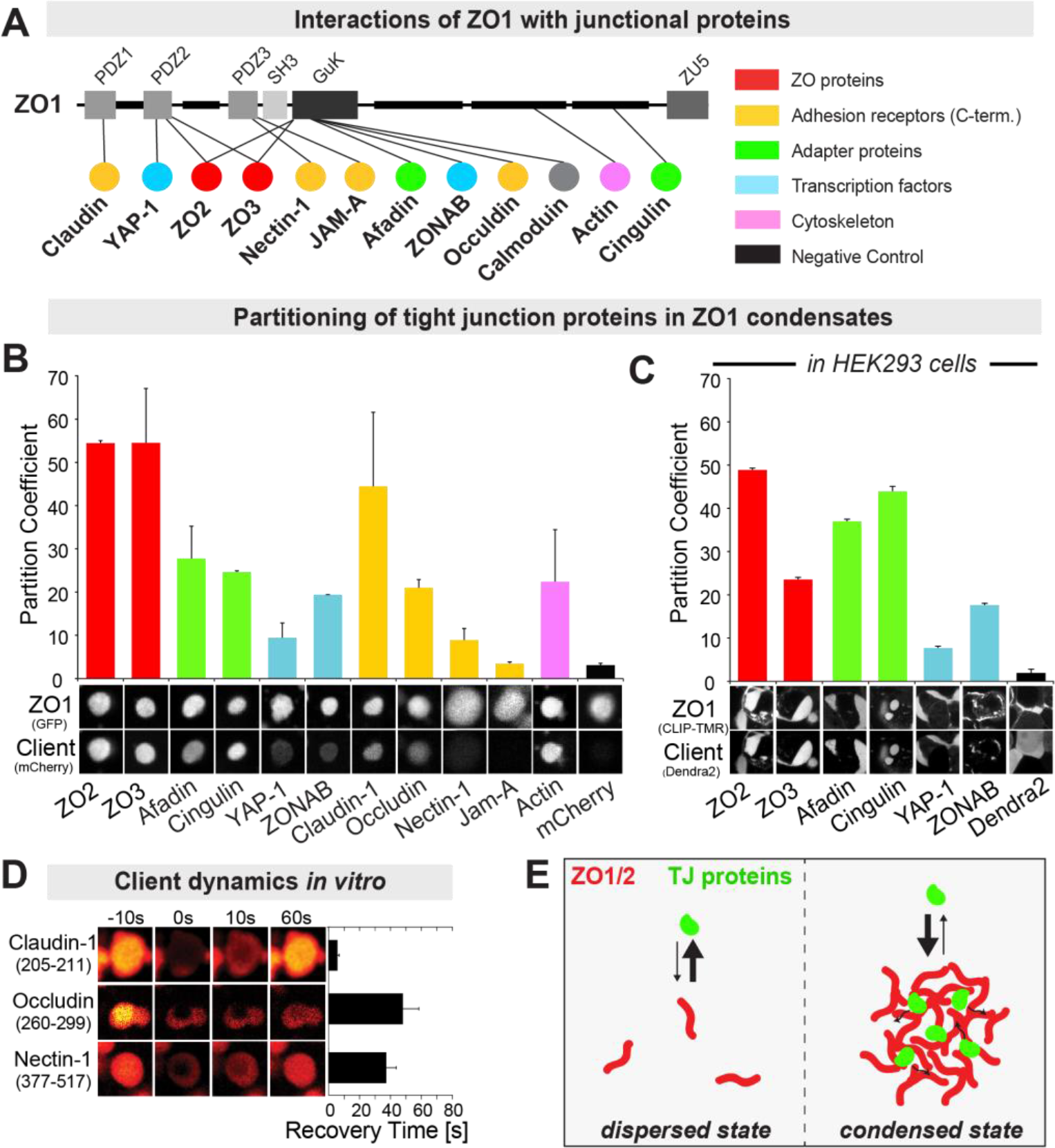
The condensed phase of ZO1 selectively sequesters tight junction proteins. (A) Scheme of known interaction sites of ZO1 with tight junction proteins. (B) *In vitro* partition assay of tight junction proteins (clients) into phase separated ZO1 compartments. The majority of clients partitioned strongly into ZO1 compartments, while the control protein (mCherry) did not. The C-terminal cytosolic fragments of TJ adhesion receptors (yellow) showed differential partitioning into the ZO1 phase, with claudin-1 having the highest and Jam-A the lowest affinity (n > 10 droplets, ± SD). (C) Partitioning of soluble tight junction proteins labelled with Dendra2 into condensed ZO1-CLIP-TMR droplets in HEK293 cells. Overall, we observed a comparable partitioning of the client proteins as seen in vitro (n > 10 droplets, ± SD). (D) FRAP measurements of client proteins *in vitro* showed that interactions between ZO1 and client proteins are transient, as indicated by the fast recovery of the protein within the ZO1 compartment. (E) Scheme of local enrichment of tight junction proteins by partitioning into condensed ZO1 compartments. In the condensed state, low affinity binding of TJ proteins to ZO1/2 is sufficient for strong partitioning due to the very high local concentration of binding sites.

To test this, we expressed and purified a number of known ZO interaction proteins (clients) and measured their partitioning into the condensed phase of ZO proteins *in vitro* (Figure 5B, S4B). First, we tested the partitioning of ZO2/3 into ZO1 compartments. As already shown in Figure 4 both homologs efficiently partition into ZO1 compartments. Next, we tested partitioning of four different adhesion receptors, which have been shown to interact with ZO1/2 (Fanning and Anderson, 2009). To avoid handling full-length adhesion receptors, we expressed the soluble cytoplasmic tails of the trans-membrane proteins (C-terminal), which include the ZO1 interaction sites. The partitioning analysis showed that the C-terminal tail of one of the main adhesion receptors of mature tight junctions, claudin-1, became 44-fold enriched in the condensed ZO1 phase. Also, occludin showed highly specific partitioning (21-fold) into the ZO1 phase. Partitioning of nectin-1, which is part of the related adherens junction complex, was significantly lower (9-fold). Surprisingly, JAM-A, which is part of the tight junction, was only slightly enriched over the negative control (mCherry). The differential partitioning of adhesion receptors into the ZO1 phase reflects their binding affinities and it may help to understand how the super-molecular structure of tight junction strands is organized.

In addition to adhesion receptors, the tight junction scaffold provides a connection to the cytoskeleton and the adherens junction. This is accomplished by secondary adapter proteins. We therefore tested the partitioning of two important representatives of this class (afadin and cingulin) (Balda and Matter, 2008; Citi et al., 2012; Ooshio et al., 2010). We found that both afadin, which connects the tight junction to the adherens junction, and cingulin, which connects the tight junction to microtubules and the acto-myosin cortex, were highly enriched in the condensed phase of ZO1 (28-fold and 25-fold respectively). We also found that monomeric actin itself became strongly enriched with the ZO1 phase (22-fold).

On top of its structural function the tight junction is also implicated in gene regulation. There is evidence that transcription factors are regulated by the cell density dependent assembly of tight junctions (Spadaro et al., 2014). Both, ZO1 and ZO2 directly interact with the transcription factor ZONAB, which regulates cell proliferation, and with the transcription factor YAP, which as part of the Hippo-pathway regulates cell growth and tissue size (Oka et al., 2010; Remue et al., 2010). Here we tested partitioning of these two transcription factors into ZO1 compartments *in vitro*. We found that both ZONAB and YAP were enriched in the ZO1 compartments (20-fold, 9-fold).

To verify the *in vitro* partitioning results, we co-expressed ZO1 with the cytoplasmic client proteins in HEK293 cells and determined its partitioning in phase-separated ZO1 domains (Figure 5C). The measurements in cells showed a slight reduction of the partitioning strength compared to *in vitro* for most proteins. However, the qualitative result was similar to the *in vitro* results. All clients with strong partitioning *in vitro* were also strongly enriched in cellular ZO1 compartments.

Taken together, the client partitioning experiments demonstrated that phase separation of ZO1 into condensed compartments results in a strong local enrichment of proteins required for tight junction assembly and signalling (adhesion receptors, cytoskeleton adapters and transcription factors). In the same fashion, FRAP experiments showed that client proteins remain mobile within the ZO1 compartments and turn over rapidly with the bulk pool of the protein (Figure 4D). Hence, phase separation of ZO1 leads to partitioning of the proteins required for tight junction assembly via specific but low affinity binding to ZO1 protein-protein interaction domains (Figure 4E).

### Phase separation of ZO1 is required for assembly of a functional tight junction belt in MDCK-II cells

To test the functional relevance of ZO1 phase separation we searched for ZO1 mutations with altered phase separation properties and probed their ability to form junctional belts in MDCK-II cells depleted of endogenous ZO1 and ZO2. Fortunately, many truncation mutants of ZO1 have already been characterized in terms of their ability to bind or to assemble tight junctions (Fanning et al., 2007; Rodgers et al., 2013; Umeda et al., 2006). Strikingly, in these studies mutations in the PSG supra-domain (SH3, U5, GuK), which we identified to be required for phase separation (Figure 4), robustly inhibit junction formation. However, because the PSG region also directly binds to other tight junction proteins (Figure 5A), interpreting these phenotypes as proof for a phase separation mechanism is not straight forward. We therefore focused on the intrinsically disordered C-terminus of ZO1. In particular, the U6 domain, which has been shown to regulate tight junction formation but is not known to bind other proteins. It was previously shown that deletion of the U6 domain causes increased assembly of ectopic tight junction strands in the lateral membrane domain in MDCK-II cells (Fanning et al., 2007; Lye et al., 2010; Rodgers et al., 2013).

To test the role of the disordered C-terminus including the U6 domain on phase separation we constructed four ZO1 truncation mutants (Figure 6A). We then used our quantitative microscopy assay in HEK293 cells to determine the phase separation properties of these mutants (Figure 6B, S6A). The first surprising finding was that removing the mostly disordered C-terminus until the U6 domain (ΔC mutant) completely supressed ZO1 phase separation in HEK293 cells. Strikingly, further truncation by removing also the U6 domain (ΔU6ΔC mutant) had the opposite effect. Phase separation was significantly increased compared to full-length ZO1. Analysis of the saturation concentrations of these mutants confirmed the strong impact of the U6 domain on phase separation compared to FL-ZO1 (Figure S6A). The plot shows that the U6 domain inhibits ZO1 phase separation and its removal promotes phase separation. The U6 has previously been shown to bind to the GuK domain via electrostatic interactions (Lye et al., 2010). Our data now suggests that this U6 back-binding prevents the PSG module from multimerization and phase separation. FRAP measurements revealed that the ΔU6ΔC condensates were less dynamic compared to full-length ZO1 condensates (Figure S6A). Interestingly, expression of full-length ZO1 with only the U6 domain deleted (ΔU6 mutant) restored the liquid-like properties of the condensates. This indicates that the intrinsically disordered C-terminus beyond the U6 domain provides liquid-like properties to ZO1 condensates, potentially by acting as a spacer between the interacting N-terminal domains (Banjade et al., 2015; Harmon et al., 2017, 2018). Finally, we asked whether the C-terminal actin-binding region of ZO1 is involved in phase separation. The ΔABR mutant phase separated in HEK293 cells, however as observed for the ΔU6ΔC mutant the dynamics in the condensates were reduced compared to full-length ZO1 (Figure S6A). Taken together, our mutation analysis revealed that the intrinsically disordered C-terminus regulates phase separation of ZO1. The acidic U6 domain is a negative regulator of ZO1 phase separation. The C-terminus downstream of the U6 is required for phase separation (in case the U6 is present), potentially by releasing the U6 inhibition. The actin-binding regions seem to be not required for ZO1 phase separation but tunes its liquid-like properties in HEK293 cells.

**Figure 6.**
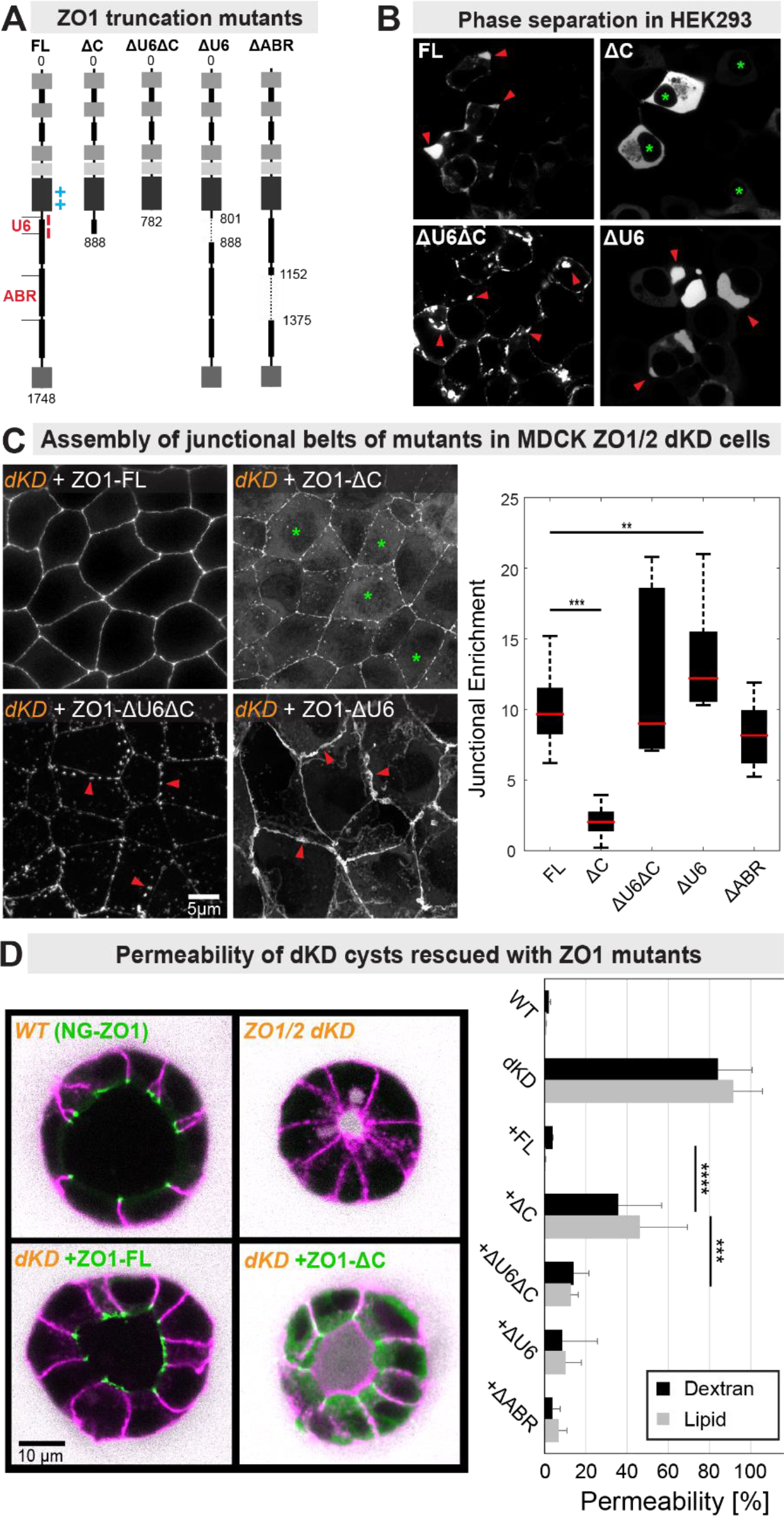
Phase separation of ZO1 is required for assembly of a functional tight junction belt in MDCK-II cells. (A) Scheme of ZO1 C-terminal truncation mutants. The positive/negative charges of the GuK and U6 domain and actin binding region (ABR) are highlighted. (B) Transient expression of ZO1 truncation mutants with N-terminal Dendra2 in HEK293. Bright condensates are indicated by red arrows. All mutants formed condensates, except the ΔC mutant, which was dispersed in the cytoplasm even at high expression levels (green stars). (C) Ability of ZO1 mutants to form continuous sub-apical belts in MDCK-II ZO1/2 dKD monolayers. All images show maximum projections of confocal stacks. Endogenous ZO1 formed continuous sub-apical belts in WT MDCK-II monolayers (Figure S6D). Knock down of ZO1 and ZO2 via CRISPR resulted depletion of ZO1/2 to levels below 4% of WT and no tight junctions were formed (Figure S6 B-D). The sub-apical ZO1 belt was restored in dKD cells by expressing ZO1-FL. The ΔC mutant did not rescue formation of a closed sub-apical belt. It was mainly localized to the cytoplasm (green stars). The ΔU6ΔC mutant formed bright membrane-bound domains but failed to assemble a continuous sub-apical belt, as seen by various gaps (red marker). The ΔU6 formed an extended sub-apical belt which often extended into the lateral membrane. We quantified the junctional enrichment of ZO1 mutants by normalizing the intensity at the junction to the intensity in the cytoplasm. The analysis confirmed that the ΔC mutant had a strongly reduced (5-times) junctional enrichment compared to ZO1-FL (n > 40 cells, ± SD). (D) Permeability of MDCK dKD cysts grown in matrigel to dextran (10k-Alexa647) and a lipid analogue (DPPE-TMR). Shown are confocal middle-planes through cysts made from WT (NG-ZO1 CRISPR KI) and ZO1/2 dKD cells that expressed Dendra2-ZO1-Mutants. In WT cysts dextran (white) did not penetrate into the lumen after 30min incubation. Also the lipid-probe (magenta) did not reach the apical side. ZO1/2 dKD cysts had severely smaller and contracted luminal shapes, which were permeable both to dextran and lipids. Expression of ZO1-FL rescued lumen shape and epithelial sealing. Expression of ZO1-ΔC partly rescued lumen shape, but epithelial permeability was significantly higher compared to WT and ZO1-FL. The other mutants are shown in figure S6E. Quantification of epithelial permeability shown on the right confirmed that the ZO1-ΔC was not able to fully rescue TJ sealing (n = 20 cysts, ± SD).

Next, we investigated the functional consequence of the mutations in terms of their ability to rescue formation of sub-apical belts in ZO1 and ZO2-depleted epithelial cells (MDCK-II ZO1/2 dKD). To deplete ZO1 and ZO2 we introduced frame-shift mutations at the respective N-termini of ZO1 and ZO2 in MDCK-II cells using CRISPR/Cas9. Immunostaining and western blot quantification confirmed that expression of endogenous ZO1 and ZO2 was reduced to less than 4% of wild type levels, respectively (Figure S6B). Measurements of trans-epithelial permeability revealed that dextran and lipid tracers had full access to the lumen of the ZO1/2 dKD cyst, while they were completely excluded in WT cells (Figure 6D). Hence, the knock-down of ZO1/2 resulted in a loss of functional tight junctions, which is in line with previous RNAi-KD experiments (Fanning et al., 2012; Rodgers et al., 2013; Umeda et al., 2006). We then used the ZO1/2 dKD cell line to perform rescue experiments with the ZO1 truncation mutants (Figure S6C). First, we confirmed that expression of full-length ZO1 resulted in formation of continuous sub-apical ZO1 belts comparable to WT cells (Figure 6C). Together with the structural rescue also the epithelial permeability was reduced to WT levels (Figure 6D). Next, we tested the localization of the ZO1 truncation mutants. To quantify the localization of ZO1 mutants with respect to ZO1-FL, we computed a junctional enrichment factor by normalizing the average intensity at the sub-apical zone to the average intensity in the cytoplasm per cell. This analysis was performed for over 40 cells from a pool of stably-transfected cells for each mutant (Figure 6C). Expression of the ΔC mutant, which did not phase separate in HEK293 cells, resulted in predominant cytoplasmic localization of the protein and only weak enrichment at the sub-apical zone. Hence, the junctional enrichment factor was significantly decreased comparted to ZO1-FL (5-fold). In contrast, the ΔU6ΔC mutant, which strongly phase separated in HEK293 cells, formed abundant ectopic ZO1 clusters and networks in the lateral membrane. However, the ectopic ΔU6ΔC clusters did in most cases not coalesce to form a continuous sub-apical belt. Interestingly, expression of the ΔU6 mutant, which contained the rest of the disordered C-terminus, resulted in formation of continuous sub-apical belts with a significantly increased expansion into the lateral membrane domain compared to ZO1-FL. As a result, the junctional enrichment factor was slightly increased compared to ZO1-FL. The expression of the ΔABR mutant, which had no strong influence on phase separation in HEK293 cells, resulted in formation of sub-apical belts comparable to ZO1-FL. Finally, the dextran and lipid permeability assay confirmed that only ZO1 mutants, with the ability to phase separate and assemble a closed sub-apical belt, were able to seal 3D cysts comparable to full-length ZO1. Notably, expression of the ZO1-ΔC mutant resulted in the highest tissue permeability, which could be significantly improved by further deletion of the U6 domain in the ΔU6ΔC mutant (Figure 6D, S6E).

Taken together, the rescue experiments revealed that the ability of ZO1 mutants to assemble continuous sub-apical belts and restore epithelial sealing strongly depended on their ability to phase separate. ZO1 mutants with decreased phase separation ability failed to assemble into continuous junctional belts. ZO1 mutants with increased phase separation formed thicker junctional belts than ZO1-FL. We, therefore, conclude that the supra-molecular assembly of ZO proteins into a dynamic compartment underlies formation of a functional tight junction belt.

## Discussion

### Phase separation of ZO proteins

We have demonstrated that all three homologs of mammalian ZO proteins can undergo spontaneous transitions into condensed liquid phases *in cellulo* and *in vitro*. Analogous to the tight junction plaque, ZO proteins are highly concentrated in the phase separated compartments and turn over rapidly with their cytoplasmic pool. The condensed ZO phase specifically sequesters and concentrates essential tight junction proteins including adhesion receptors and cytoskeletal adapters. These results support the hypothesis that phase separation of ZO protein is important for tight junction assembly. We, therefore, propose that sequestering and segregation of tight junction components to nascent cell adhesion sites is driven by a phase transition of ZO1/2 into a condensed membrane attached compartment. Local enrichment and scaffolding within the compartments may then enable polymerization of claudin and actin into the mature junctional network. We note that polymerization of claudin and actin are expected to change the material state of the ZO1 scaffold towards a gel or solid like assembly rather than a disordered protein liquid. In fact, FRAP experiments on tight junctions have shown that the dynamics of claudin receptors are very low (Shen et al., 2008), which indicates solid like material properties of the trans-membrane proteins. In addition, also ZO1 has a 30% fraction that is not dynamic at the junction. Presumably, this is the fraction that is directly bound to the polymerized claudin strands.

We have identified that the protein-protein interactions of the conserved PSG supra-domain of ZO proteins are essential for phase separation (Figure 4). This is in agreement with previous ideas that the PSG domain of MAGUK proteins may assemble into polymers, for example by domain-swapping (Fanning and Anderson, 2009; Pan et al., 2011; Umeda et al., 2006; Ye et al., 2018). Further support for this model comes from our observation that the phase separation affinity of ZO3 is significantly higher than ZO1/2 (Figure 3D). The increase phase separation affinity correlates with a more open conformation of the PSG domain of ZO3 compared to ZO1 (Lye et al., 2010), which may facilitate polymerization and hence phase separation. A more closed packing of PSG domains in other MAGUKs such as PSD95 may also explain why these scaffolding proteins require interactions with additional proteins to assemble into scaffolds (Zeng et al., 2016). Importantly, we found that the U6 domain of ZO1, which has been shown to regulate tight junction formation by binding to the PSG domain (Fanning et al., 2007; Lye et al., 2010; Rodgers et al., 2013), inhibits the phase separation ability of ZO1 and dramatically reduces junction formation (Figure 6). This finding suggests a mechanism for how phase separation of ZO proteins can be actively controlled in space and time.

### Implications of phase separation mediated tight junction assembly

Assembly of tight junctions requires a mechanism to accumulate the necessary proteins at the sub-apical region and facilitate formation of a continuous belt by polymerization of claudin receptors and maybe actin filaments. Based on the large body of tight junction studies and our new data, we postulate that the assembly of the mature junctional complex involves three steps (Figure 7).

**7.**
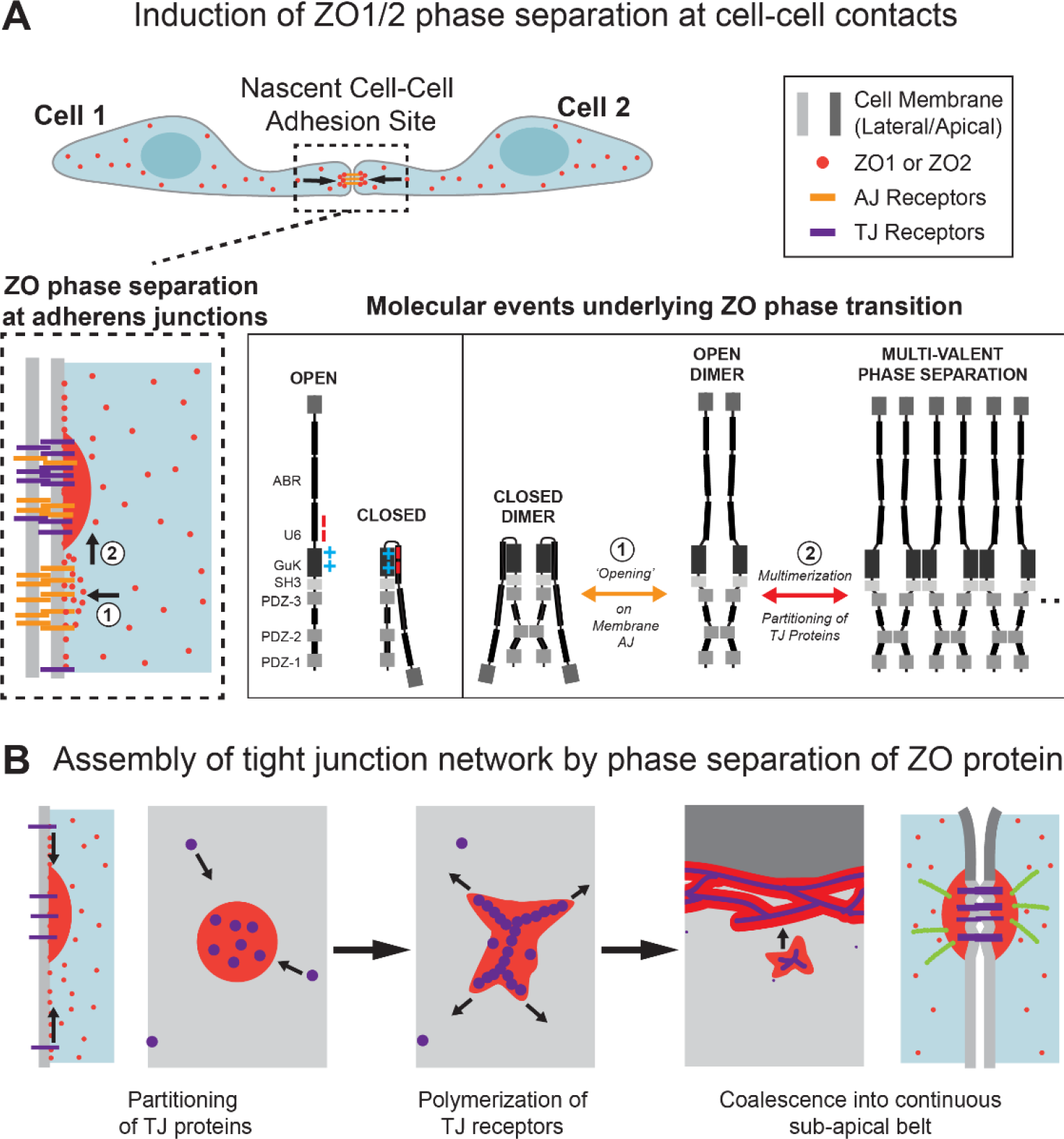
Model of tight junction formation by phase separation of ZO1/2. (A) Potential induction of ZO phase separation at nascent cell-cell contact sites. ZO is recruited to early adhesion sites via adherens junction (AJ) receptors and adaptor proteins. Membrane recruitment may be sufficient to cross the concentration threshold for phase separation. However, our experiments suggest that ZO1 is self-inhibited by its U6 domain. We refer to the inhibited state as the closed state, because the U6 domain is folded back. In order to promote tight junction strand assembly ZO1 needs to be opened. Opening releases the self-inhibition and promotes phase separation via multimerization of the PSG domain. The mechanism how ZO1 can be opened could be via de-/phosphorylation by junction-specific phosphatases/kinases or via mechanical forces applied to the C-terminus by acto-myosin. (B) Formation of ZO dense compartments causes partitioning of tight junction-specific proteins including claudin receptors. The local accumulation and scaffolding of claudins may be sufficient to trigger polymerization and strand formation. Due to the fluid-like nature of the ZO1/2 compartments coalescence of multiple growing compartments into a continuous belt is facilitated.

First, ZO1/2 are recruited to nascent adhesion sites by the formation of adherens junctions (Ando-Akatsuka et al., 1999; Yonemura et al., 1995). Binding of ZO proteins to adherens junctions will increase its local concentration, which may already be sufficient to cross its phase separation threshold. However, our results on the inhibition of ZO1 phase separation by the U6 domain, suggest that ZO proteins need to be actively released from auto-inhibition to promote phase separation (Figure 7A). This active process could involve binding of another protein, de-/phosphorylation, or even require mechanical force applied by the cortex (Spadaro et al., 2017). Evidence for an acto-myosin-dependent clustering of ZO proteins has been found during zebrafish embryogenesis (Schwayer and Heisenberg et al. unpublished data)

Second, our experiments have shown that phase separation of ZO1/2 produces compartments that specifically sequester tight junction proteins and locally concentrate them up to 40-fold over the bulk phase (Figure 7B). The dynamic partitioning of junctional proteins is a consequence of the relatively low affinity interactions of ZO1 with tight junction proteins in combination with the high number of local binding sites within the phase separated scaffold. Addtionally, the differential affinity of the ZO1 scaffold for adhesion receptors (claudin > occludin > nectin > jam-a) suggest a hierarchical assembly process. However, additional information on the stoichiometry of the components in cells will be required to make further predictions about this.

Third, we propose that the partitioning of junctional proteins is sufficient to drive polymerization reactions and facilitate tight junction strand formation. In accordance with this idea, claudin receptors do not polymerize into a continuous network in epithelial cells in the absence of ZO1/2 (Fanning and Anderson, 2009; Umeda et al., 2006). However, heterologous overexpression of claudin receptors in other cell types induces spontaneous formation of typical tight junction strands at cell-cell contacts independent of ZO proteins (Furuse et al., 1998). Therefore, nucleation of claudin strands seems to require a threshold concentration of claudin monomers at the plasma membrane. We propose that the polymerisation concentration of endogenous claudin receptors in epithelial cells is reached solely when the receptors are locally concentrated and scaffolded by phase separated ZO compartments.

Altogether, based on our experiments we propose that tight junction assembly involves a transition of ZO proteins into a condensed membrane bound phase, which sequesters junction-specific components and nucleates claudin and actin polymerization. Our data suggest that the ZO phase transition into a condensed state is controlled by intra-molecular inhibition and requires an active process to release the inhibition. This step could involve phosphorylation or mechanical opening at adherens junctions. We note that the mature junctional complex, with its layered molecular organization, shows features of both a viscous liquid as well as an ordered crystal. Our work opens the door to develop a mesoscale understanding that captures both of these properties and can serve as a template to reconstitute the supra-molecular organization of the tight junction complex.

## Acknowledgments

We thank the Protein Expression & Purification, Genome Engineering, Cell Technologies and Mass Spectrometry facilities at the MPI-CBG; Christoph A. Weber and Lars Hubatsch for help with the FRAP data; Titus Franzmann for help with phase separation assays; This work was funded by the Max Planck Society, by the Deutsche Forschungsgemeinschaft (DFG, German Research Foundation) - Project Number 112927078 - TRR 83 and Project Number 4191381812. KPG was supported by an HFSP research grant.

## Author Contributions

A.H. and O.B. conceived the project and analysed the data. K.P-G, O.B. and A.H. wrote the manuscript. O.B. performed all experiments, except those specifically attributed to other authors. R.M. performed Crispr-knock downs and epithelial permeability experiments. K.P-G. helped in protein purification and protein quality control. C.M-L did the cell culture work and created all stable cell lines.

---

**Figure S1.**
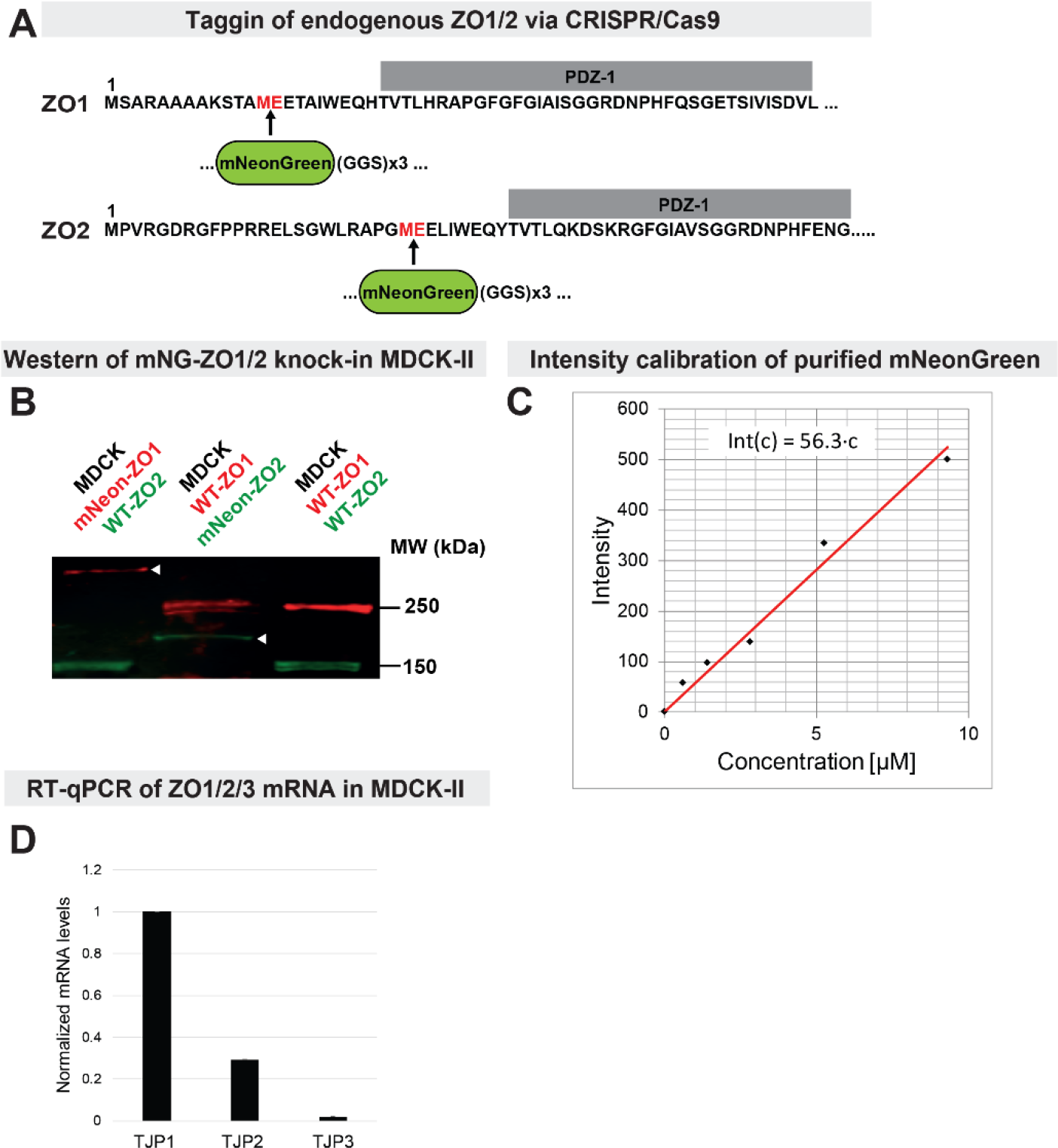
Fluorescent tagging and quantification of endogenous ZO1/2. (A) CRISPR/Cas9-mediated insertion of mNeonGreen including a GGS linker at the indicated position before the PDZ1 domain of ZO1/2. (B) Western blot against ZO1 (red) and ZO2 (green) in WT MDCK-II and CRISPR/Cas9-Mediated fluorescent tagged MDCK-II cells. White arrows indicate the shift in molecular weight of ZO1 and ZO2 due to the mNeon insertion. (C) Intensity versus concentration calibration curve of purified mNeon with the same settings used for imaging cells in Figure 1C. (D) Real-time PCR of ZO1/2/3 (TJP1/2/3) in WT MDCK-II cells. The results are normalized to ZO1 levels. The ratios between ZO1 and ZO2 match the quantification of protein levels determined by imaging and FCS (Figure 1 C,D,G).

**Figure S2.**
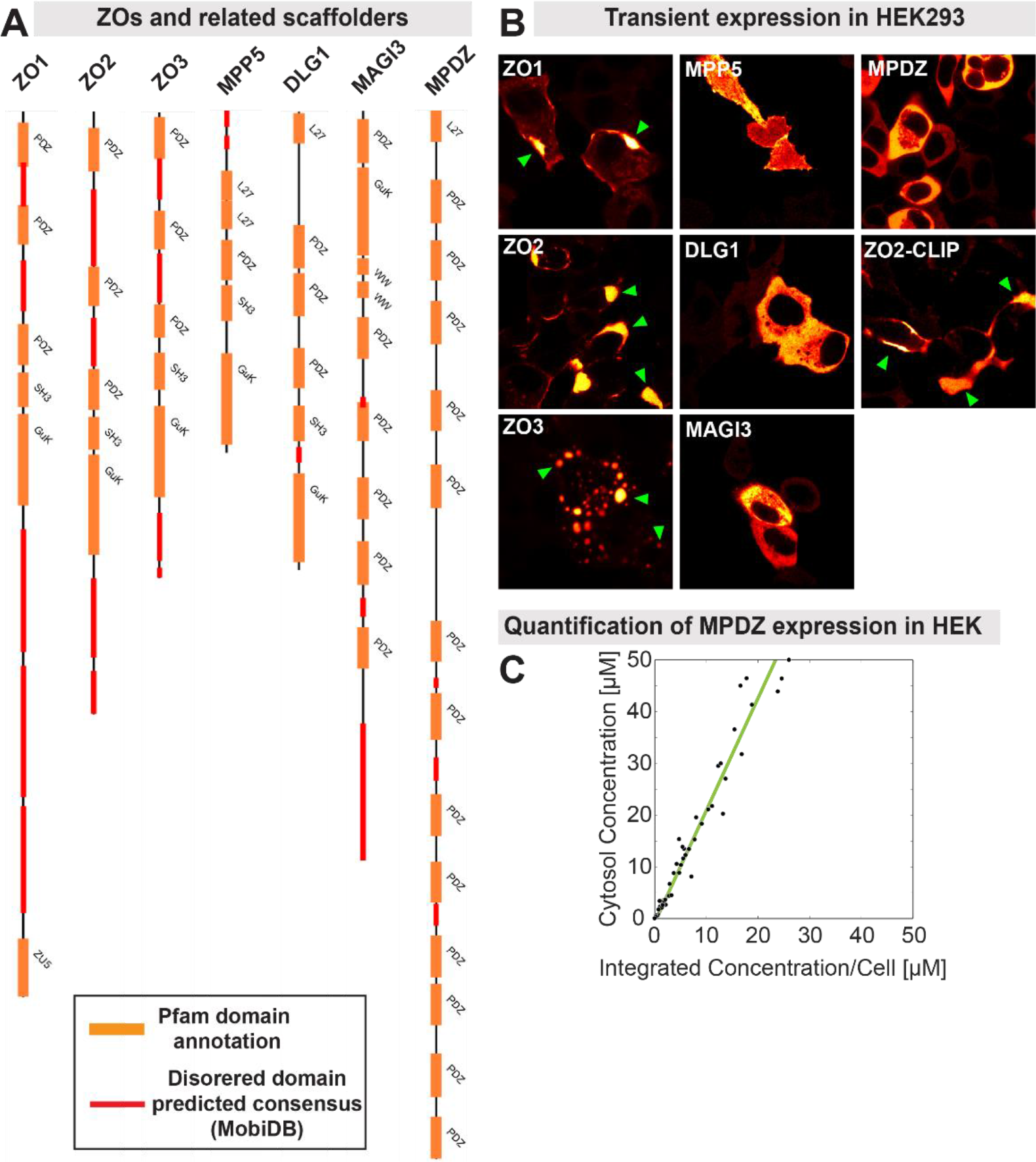
Transient expression of multi-domain scaffolding proteins in HEK293. (A) Domain structure of ZO proteins and four related scaffolding proteins. Predicted intrinsically disordered regions are shown in red. (B) Representative images of the proteins expressed with N-terminal Dendra2 tags in HEK293. Formation of condensed domains is indicated with green arrows. Only ZO proteins were able to form bright condensates. Exchanging the tag on ZO2 to CLIP and staining with TMR resulted in formation of similar condensates as for Dendra2-ZO2, indicating no influence of the tag. (C) In line with the visual impression, quantification of MPDZ expression level versus cytoplasmic concentration showed a linear relation and no signs of saturation.

**Figure S4.**
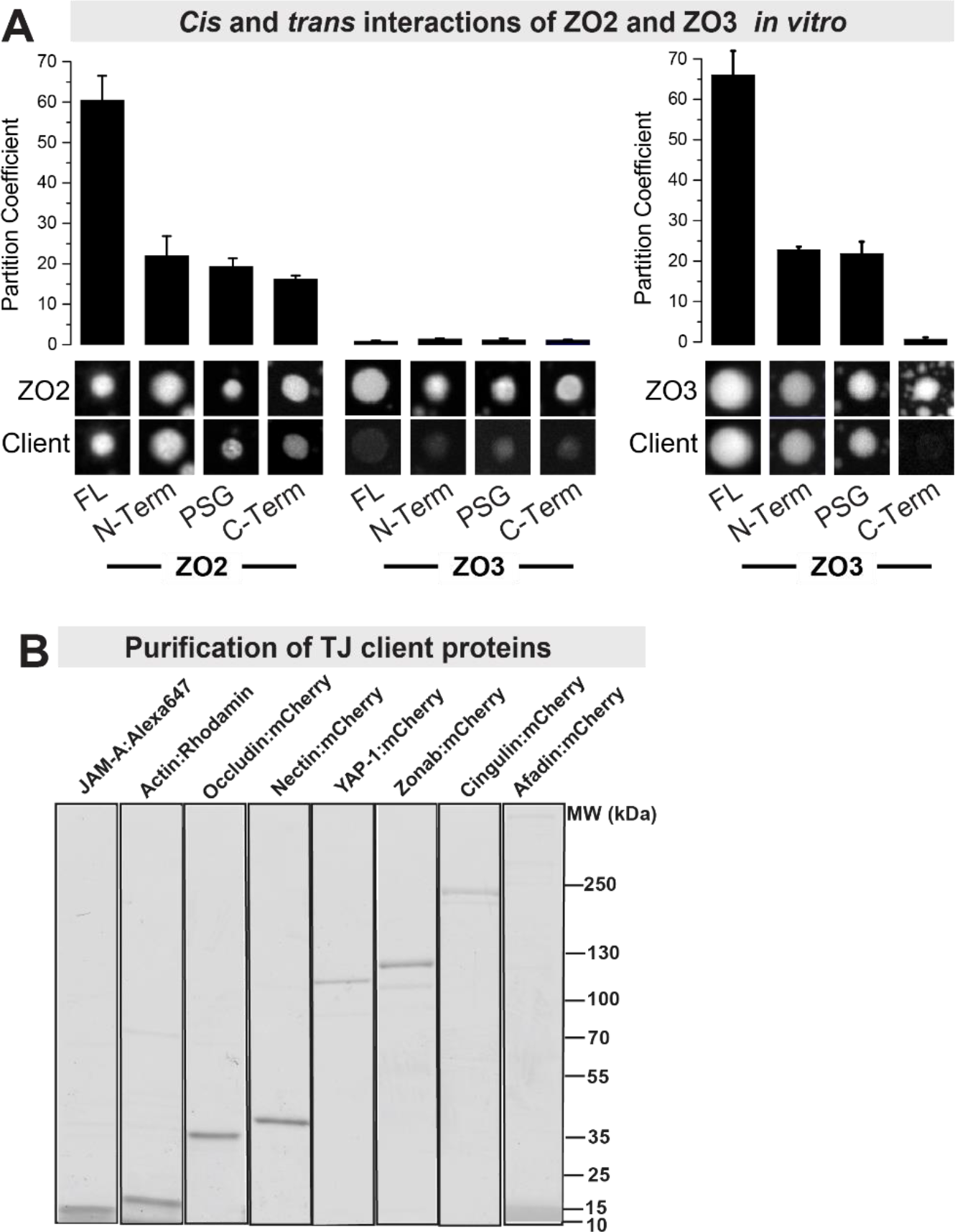
Partition assay for ZO2 and ZO3. **(A)** Partitioning assay to determine protein-protein interactions of phase separated full-length ZO2 or ZO3 with fragments of ZO2 and ZO3 (Clients). Partitioning was determined by computing the ratio of fluorescence inside to outside of the droplet for the client proteins. The results show that ZO2 and ZO3 show similar *cis*-interaction with their N-terminal, PSG and C-terminal fragment as ZO1, except that the ZO3 c-terminus did not interact with ZO3-FL. Interestingly, ZO2 and ZO3 did not interact with each other in *trans*. (B) Purification of tight junction client proteins for the parititioning assay in Figure 5.

**Figure S6.**
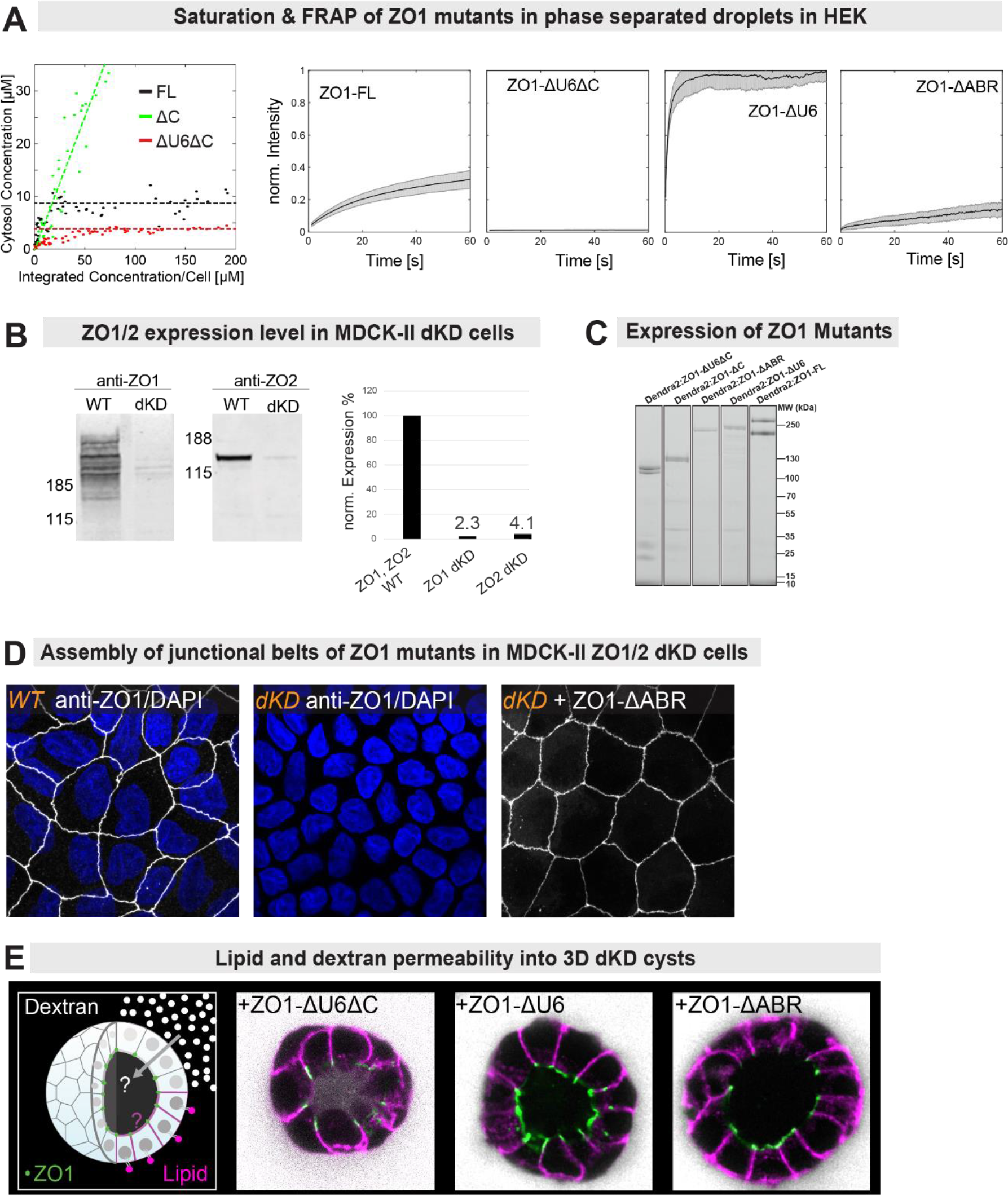
Quantification of ZO1 truncation mutants in HEK and MDCK ZO1/2 dKO cells. **(A)** Saturation concentration for phase separation of the ΔC and ΔU6ΔC mutant compared to ZO1-FL in HEK293. Full length (FL) ZO1 saturated and phase separated around 10 μM. The ΔC mutant did not show any sign of saturation even at high expression levels. The ΔU6ΔC phase separated at even lower saturation than ZO1-FL (n > 60 cells). FRAP curves for ZO1 truncation mutants in the condensed phase in HEK293. The ΔU6ΔC and to a lesser extend the ΔABR mutants showed significant decrease in the recovery of the beached condensates, indicating a more solid-like material state. (B) Western blot quantification of ZO1 and ZO2 protein levels after CRISPR/Cas9 double knock-down (dKD) of ZO1/2. The analysis showed a strong decrease > 95% of ZO1 and ZO2 protein levels. (C) In-gel-fluorescence of MDCK-II dKD extracts showing the correct expression of Dendra2-tagged ZO1 truncation mutants. (D) Expression and localization of endogenous ZO1 in WT MDCK-II monolayers. In MDCK-II ZO1/2 dKD monolayers ZO1 does not form a sub-apical belt. Expression of the ZO1 ΔABR mutant rescued belt formation. (E) Trans-epithelial permeability of ZO1/2 dKD cells was tested by measuring permeability of fluorescent dextran (10kD) and a fluorescent lipid analogue (2kD) into the lumen / apical site of 4-days old MDCK cysts cultured in Matrigel. ZO1/2 dKD expressing ZO1-ΔU6ΔC did not fully rescue the barrier and fence function of the tight junction. Expression of ZO1-ΔU6 and ZO1-ΔABR largely rescued junctional sealing.

## Material and Methods

### KEY RESOURCES TABLE

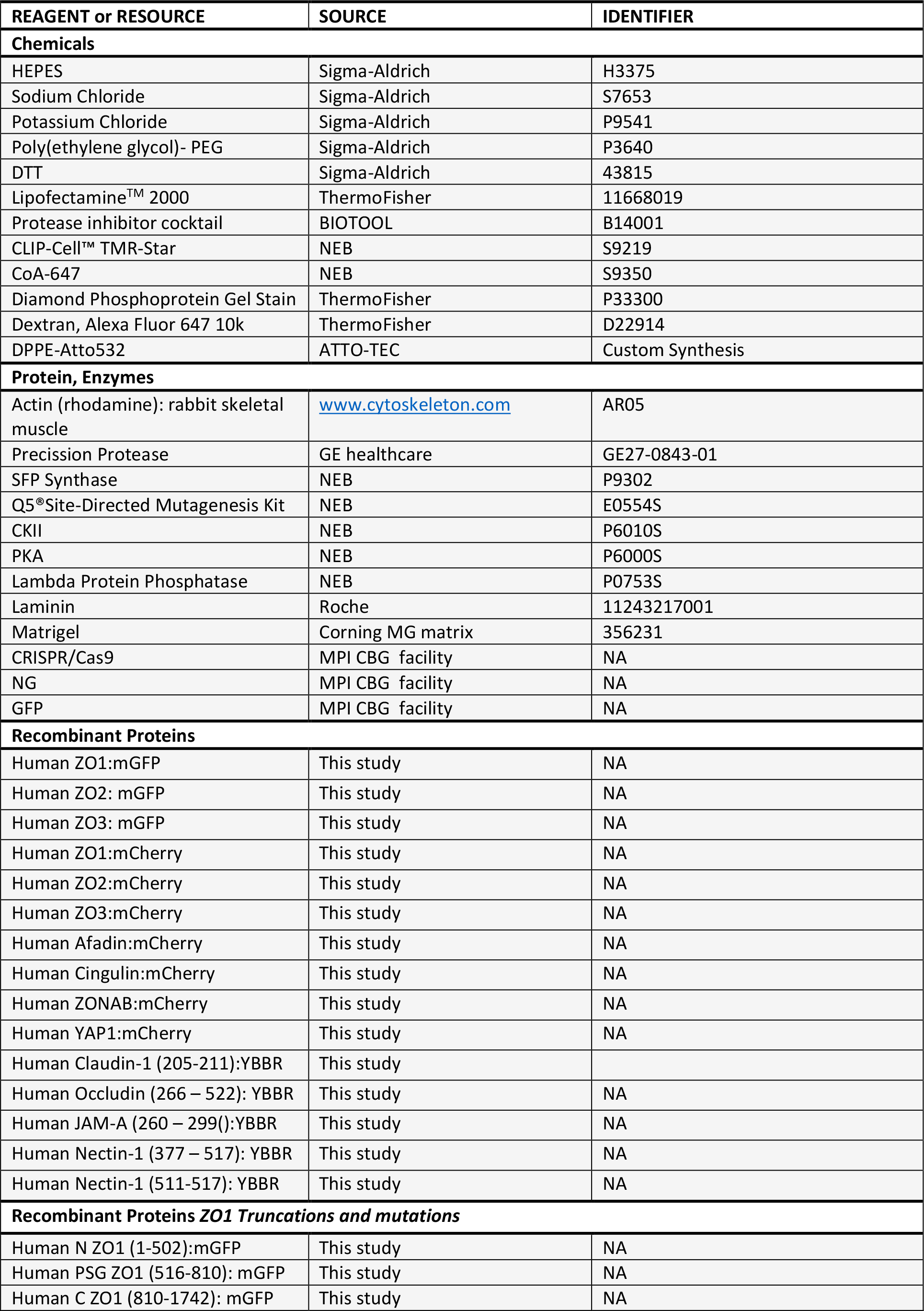

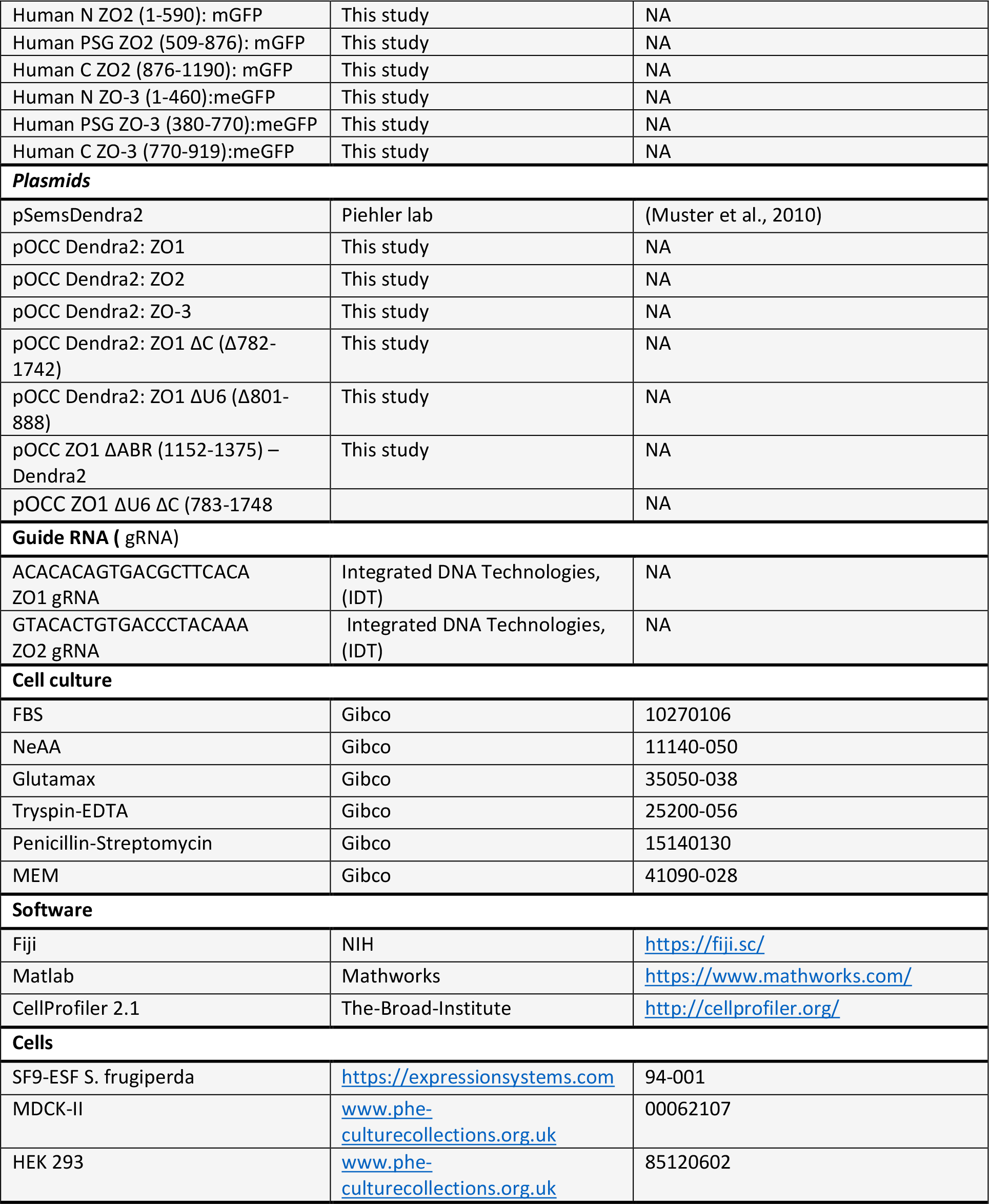

### CONTACT FOR REAGENT AND RESOURCE SHARING

Further information and requests for resources and reagents should be directed to and will be fulfilled by the Lead Contact, Alf Honigmann

### QUANTIFICATION AND STATISTICAL ANALYSIS

Images were analyzed with FIJI (https://fiji.sc/) and MATLAB (Mathworks). All data are expressed as the mean ± the standard deviation (SD), mean ± the standard error of the mean (SEM), or mean ± 95% confidence intervals as stated in the figure legends and results. The value of n and what n represents (e.g., number of images, condensates or experimental replicates) is stated in figure legends and results. Two-tailed Student’s t tests were used for normally distributed data, and Wilcoxon Rank-Sum tests were used for non-normally distributed data. A Pearson’s Chi-square test was used to determine if data were distributed normally.

#### Cloning

The proteins were codon optimized for eukaryotic expression and de novo synthesized by GenScript.

##### Insect expression system and vectors

The synthesized genes were cloned via Not-I and Asc-I cutting sites into in house designed baculoviral expression plasmids (pOCC series) for expression

| Plasmid name | Gene | N-Term Tag | C-Term Tag | Origin |
| --- | --- | --- | --- | --- |
| AH Z01-GFP | Human TJP1 | HIS6-MBP-3C | TEV-monoGFP | This study |
| AH Z01-mCherry | Human TJP1 | MBP-3C | mCherry-3C- His6 | This study |
| AH Z01 N-mCherry | Human TJP1 | MBP-3C | mCherry-3C- His6 | This study |
| AH Z01 PSG-mCherry | Human TJP1 | MBP-3C | mCherry-3C- His6 | This study |
| AH Z01 C-mCherry | Human TJP1 | MBP-3C | mCherry-3C- His6 | This study |
| AH-Z02-GFP | Human TJP2 | HIS6-MBP-3C | TEV-monoGFP | This study |
| AH-Z02-mCherry | Human TJP2 | MBP-3C | mCherry-3C- His6 | This study |
| AH Z02 N-mCherry | Human TJP2 | MBP-3C | mCherry-3C- His6 | This study |
| AH Z02 PSG-mCherry | Human TJP2 | MBP-3C | mCherry-3C- His6 | This study |
| AH Z02 C-mCherry | Human TJP2 | MBP-3C | mCherry-3C- His6 | This study |
| AH Z03-GFP | Human TJP3 | HIS6-MBP-3C | TEV-monoGFP | This study |
| AH Z03-mCherry | Human TJP3 | MBP-3C | mCherry-3C- His6 | This study |
| AH Z03 N-mCherry | Human TJP3 | MBP-3C | mCherry-3C- His6 | This study |
| AH Z03 PSG-mCherry | Human TJP3 | MBP-3C | mCherry-3C- His6 | This study |
| AH Z03 C-mCherry | Human TJP3 | MBP-3C | mCherry-3C- His6 | This study |
| AH cingulin –mCherry | Human Cingulin | MBP-3C | mCherry-3C- His6 | This study |
| AH Afadin –mCherry | Human Afadin | MBP-3C | mCherry-3C- His6 | This study |
| AH Zonab –mCherry | Human ZONAB | MBP-3C | mCherry-3C- His6 | This study |
| AH Yap-1 –mCherry | Human Yap-1 | MBP-3C | mCherry-3C- His6 | This study |

##### Mammalian expression system and vectors

The synthesized genes were cloned via Not-I and Asc-I cutting sites into in house designed mammalian expression plasmids (pOCC series) with N-terminal Dendra-2, selection marker against Neomycin-Geneticin and a CMV promotor. The used constructs are summarized in the following table:

| Plasmid name | Gene | N-Term Tag | C-Term Tag | Origin |
| --- | --- | --- | --- | --- |
| AH 62 Z01 –Dendra2 | TJP1 | Dendra2 |  | This study |
| AH 148 Z01 ΔU6ΔC – Dendra2 Z01 ($\Delta$ 782-1742) | TJP1 | Dendra2 | | This study |
| AH 176 Z01 ΔU6 (801-888) –Dendra2 | TJP1 | Dendra2 | | This study |
| AH 174 Z01 ΔC (1-888) –Dendra2 | TJP1 | Dendra2 | | This study |
| AH 175 Z01 ΔABR (1152-1375) –Dendra2 | TJP1 | Dendra2 | | This study |
| AH 172 Z01 ΔU5 – Dendra2 | TJP1 | Dendra2 | | This study |
| **AH 66** ZO2 –Dendra2 | TJP2 | Dendra2 |  | This study |
| **AH 67** ZO3 –Dendra2 | TJP3 | Dendra2 |  | This study |
| **AH 127** cingulin – Dendra2 | Cingulin | Dendra2 |  | This study |
| **AH 131** Afadin – Dendra2 | Afadin | Dendra2 |  | This study |
| **AH 128** Zonab – Dendra2 | Zonab | Dendra2 |  | This study |
| **AH 130** Yap-1 – Dendra2 | Yap-1 | Dendra2 |  | This study |
| **AH 112** ZO1 –Clip | TJP1 | CLIP |  | This study |
| pSems Dendra2 | Dendra2 | Dendra2 |  | (Muster et al., 2010) |

#### Protein Expression and Purification

##### Protein expression and purification of scaffolding proteins

We used insect cells to recombinantly express proteins using the baculovirus expression system. SF9-ESF *S. frugiperda* insect cells transfected with the respective protein were grown at 27°C in ESF 921 insect cell culture medium supplemented with 2% Fetal Bovine Serum. After 72 h the Cells were collected, washed, and resuspended in harvest buffer (50 mM Tris HCl, pH 7.4, 150 mM NaCl, 30 mM imidazole, 1% glycerol) + protease inhibitors (1 mM PMSF, 100 mM AEBSF, 0.08 mM Aprotinin, 5 mM Bestatin, 1.5 mM E-64, 2 mM Leupeptin, 1 mM Pepstatin A) and frozen in liquid nitrogen. All following steps were carried out 4°C.

Cells were opened by using a dounce homogenizer. Lysates were clarified by centrifugation for at 75000 × g 25 minutes. Afterwards, the protein of interest was enriched via metal ion affinity chromatography (Ni-NTA) resin (IMAC, 5ml HiTrap Chelating, GE Healthcare). After 10 column washes with (20 mM HEPES, pH 8, 500 mM NaCl, 40 mM imidazole, 1mM DTT) the protein was eluted with 500 mM imidazole. His and MBP affinity tags were cleaved off using a histidine tagged precission protease (in-house) overnight. Finally, size exclusion chromatography was performed with Superose 6 column. Truncated mutant proteins were purified by Superdex 200 increase 10/300 GL column (GE). Both columns were equilibrated with storage buffer (20 mM HEPES, pH 8.0, 500 mM NaCl) and were connected to an Akta Ettan FPLC system (GE). The proteins were frozen in liquid nitrogen and stored at −80°C in 20 mM HEPES, pH 8.0, 500 mM NaCl, 1 mM DTT and 5% Glycerol at −80°C.

##### Protein expression and purification of C-terminal domains of Occludin, JAM-A and Nectin-1

The C-terminal domains of Occludin, JAM-A, Nectin-1 were expressed in *E.Coli* (BL21(DE3)) induced with 0.5 mM IPTG for 16 hr at 20°C. Cell pellets were frozen in liquid nitrogen. Re-suspended in lysis buffer (20 nM HEPES, pH 7.2, 150 mM NaCl, 10 mM imidazole, 0.1 % Triton-X 100) supplemented with protease inhibitors. Cells were lysed by sonication then clarified by centrifugation for 25 minutes, 75,000 x g at 4 C. The proteins were the purified via His-Hi TRAP and Superdex 200 as described above. The C-terminal peptides of claudin-1 and nectin-1 were synthesised via peptide synthesis by the technology platform Dresden.

Purified C-termini of Claudin-1, Occludin or JAM-A or Nectin-1 contained an YBBR-tag which was labelled with CoA-DY647 by using the PPTase Sfp as described previously (Yin et al., 2005). 10μM of the protein and 1.5 times excess of the CoA 647 substrate were mixed and incubated for 1h at room temperature. After 1 hour the excess dye was removed via PD-10 desalting column (GE Healthcare) equilibrated with the storage buffer (20 mM HEPES, pH 7.4, 500 mM NaCl). The labelled proteins were eluted from the column and stored at − 80°C.

##### Immunoblotting and in-gel fluorescence

Total cell lysates were obtained in lysate buffer (10% glycerol, 2% SDS, 1mM DTT, 1x Protease Inhibitor cocktail, 40 mM Tris-HCl, pH 7.6) were incubated with of 250 U of benzonase shaking 15 min at 37 C. Then 1mM EDTA, 1mM EGTA was added. Protein loadings were normalized by BCA. The SDS PAGE 4-20% Tris-Glycine Novex Gels were run at 90 mV for 3 hours. Transfer was done in a iBlot2 system onto a nitrocellulose membrane (10 min, 20V). To detect protein levels by inmmunoblotting a iBind system was used and the following antibodies were used: ZO1 mouse monoclonal (1:750, Invitrogen, 33-9100), ZO2 polyclonal rabbit (1:750, Cell signaling, 2847S). As secondary antibodies used IRDye-800CW goat anti-mouse (1:4000, LI-COR, 925-32210) and IRDye-680CW Goat anti-rabbit (925-68021, LI-COR). Membranes were scaned by LI-COR Odyssey system at 700 nm and 800 nm. In-Gel fluorescence of the tagged proteins (Dendra, mNeon, GFP, mCherry) was done in a Thyphoon FLA 7000 scan system (GE). Equal concentration of pure protein (20ug) or lysates (100ug) were loaded onto a 1D SDS-PAGE and the gels were further stainned with Comassine and scan using a LI-COR system.

##### De-/Phosphorylation assay of ZOs

ZO1 was phosphorylated *in vitro* using CKII kinase. ZO1 was incubated with 2.500 U CKII in 20 M HEPES pH 7.5, 1 mM DTT, 10 mM MgCl_2_, 200μM ATP and 500mM NaCl for 1 hour at 30°C. ZO1 was dephosphorylated by using lambda protein phosphatase(PP1). ZO1 was incubated with 1000U of PP1 in 20 mM HEPES pH 7.5, 1 mM DTT, 1mM MnSO_4_, and 500mM NaCl for 1hour at 30°C. Phosphorylation and de-phosphorylation of the ZO proteins was validated by 1D SDS PAGE after staining for phosphorylation using diamond phosphoprotein gel stain. Phosphorylation sites of the ZO1 were further analysed by mass spectrometry on a HF Quadrupole Orbitrap Hybrid Mass Spectrometer.

##### Phase separation and client partitioning assay in vitro

Purified ZO1/2/3 and their respective mutants were diluted from storage buffer into phase separation buffer (20 mM HEPES, pH 7.5, 150 mM KCL and 2% PEG(20k)). Titration of the final protein concentration from 0.1 to 10 μM allowed us to estimate the saturation concentration of the respective protein. Formation and fusion of liquid droplets was observed by bright field and fluorescence microscopy using a 60x water objective. For partitioning analysis, ZO proteins, labelled with GFP, were mixed with the respective interactions partner, labelled with mCherry were mixed in a molar ratio 2:1 and diluted in phase separation buffer. The partitioning was quantified by measuring the fluorescence of the client proteins inside the droplets and normalizing this to the fluorescence outside.

##### Confocal microscopy Imaging

All microscopy measurements (FRAP, FCS, imaging) were performed on a confocal microscope (Abberior Instruments, Göttingen, Germany) with pulsed laser excitation (490 nm, 560 nm, 640 nm, 40MHz).

##### Fluorescence recovery after photobleaching (FRAP)

*In vivo* experiments performed in (MDCK-II and HEK293) for bleaching the tight junction were carried out with the following settings. A region of 10 × 10 μm (pixel size 100 nm) was imaged for 3 frames, a 5 × 5 μm rectangular region of interest (ROI) (for *in vitro*) or a 3 × 0.5μm rectangular region of interest (ROI) (for bleaching the tight junction) was scanned by the 405 nm and the 488 nm lasers at 5 μW and 20 μW, respectively. Fluorescence recovery of meGFP or Dendra2.0 was monitored for 1-20 min with a time resolution of 11s. Cell movements during the recovery were corrected by registration of all frames to the first frame using the plugin StackReg in FIJI. For FRAP on the condensates recovery was measured with a frame rate of 5s was. Recovery data was background corrected and normalized to the ROI intensity prior to bleaching. A reference ROI outside the bleached area was processed in the same way. To correct for photobleaching during the measurement. FRAP traces were evaluated and fitted in MATLAB.

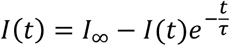

##### Fluorescence correlation spectroscopy (FCS)

mNeonGreen was excited with a 490 nm pulsed laser (40 MHz, laser power 10 μW at the back aperture of an Olympus 60x NA = 1.2 (UPLSAPOXW) water objective). Fluorescence fluctuations were recorded with a time resolution of 500 ns for 45 s. Auto-correlation of the photon traces was performed in MATLAB using a multiple tau correlator. The resulting correlation curves were fitted according to the standard 3D diffusion model including one triplet component using MATLAB (Elson, 2011). Focal aspect ratio, triplet time and triplet fraction were determined by measuring purified meGFP and NG in solution and were subsequently fixed when fitting all subsequent data. Oligomer state of the proteins was estimated by calculating the counts per molecule: *CPM = <I>/N*, with *<I>* being the mean intensity and *N* the mean particle number obtain by the FCS fit. Normalizing the CPM to the CPM of free NG expressed in the same MDCK-II cell line yielded the average number of NeonGreen per particle, i.e. the oligomeric state.

##### Absolute quantification of protein concentrations

The absolute concentration of the mNeonGreen and Dendra2 tagged proteins (ZO1 and ZO2) in MDCK cells was estimated by fluorescence microscopy. First, we calibrated the intensity of NeonGreen on our microscope as a function of laser power and protein concentration using purified NeonGreen. The conversion factor was obtained by linear fitting of the mean pixel intensity values (I) against the NeonGreen concentration (45 ± 0.5 [counts/ μM]). For the Dendra2-tagged proteins we used purified mEGFP to calibrate the conversion factor and assumed that Dendra2 brightness is 0.67 of meGFP (Gurskaya et al., 2006).

##### Determination of the saturation concentration in HEK293 cells

To determine the saturation concentration for phase separation we transiently expressed Dendra2 tagged proteins in HEK293 cells. We took 3D confocal stacks of over 100 cells 24h after transfection. To determine the cytoplasmic concentration of each cell, we measured the mean intensity in a cytoplasmic region in the mid-plane of the confocal stack and used the conversion factor for Dendra2 to calculate the protein concentration. To estimate the total concentration of Dendra2 of the same cell (including the phase separated droplets) we summed up all the pixels within the cell by using a sum-projection and a hand-drawn region of interest in FIJI. We then plotted the total expression of the protein per cell against the cytoplasmic concentration. For control proteins like Dendra2 this plot was linear as expected. In case of formation of condensates, the cytoplasmic concentration saturated at characteristic values. The saturation concentration was estimated by calculating the mean of all concentrations in the saturated plateau.

##### Determination of the tight junction enrichment factor in MDCK-II cells

To determine the tight junction enrichment factor, we measured via FiJi the intensity in the cytosol and on the junction. For this we used maximum projections of the 3D confocal stack and measuring the mean intensity in the cytosol and the mean intensity on the tight junction. The enrichment factor was then calculated *R* = (*I*_*junction*_ − *I*_*cytosol*_) * *I*_*cytosol*_.

#### Cell culture and Genetic Engineering

##### HEK293 and MDCK-II cell culture

HEK 293 and MDCK-II cells were cultivated at 37°C and 5% CO2 with DMEM supplemented with 5% FBS, 1% non-essential amino acids and 1% 2(4-(2-hydroxyethyl)-1-piperazineethanesulfonic acid (HEPES) buffer without addition of antibiotics. Transgenic cell lines were grown in presence of geneticin. Transient transfection of HEK293 cells was carried out using Lipofectamine2000 (ThermoFisher) after cells reach a confluency of 70%.

##### CRISPR/Cas9 knock-in and knock-down in MDCK-II cells

In order to knock-down ZO1 and ZO2 in MDCK-II frame-shift mutations were introduced at the N-termini by CRISPR/Cas9. The following guides were used for ZO1 ACACACAGTGACGCTTCACA and ZO2 GTACACTGTGACCCTACAAA. Selected DNA oligos and their trans-encoded RNA (TRCR) were purchased from Integrated DNA Technologies. The gRNA was annealed for 1h at room temperature with CRISPR protein and its trans-encoded RNA finally gene rate the riboprotein complex. The complex was incubated with homemade purified Cas9. Electroporation of each complex was performed in 300.000 cells (Invitrogen NEON electroporation machine and kit, 2 pulses, 20 ns, 1200 V). Cells were sorted after 48h by FACS (fluorescence activated cell sorting) and grown clonally. The genomic sequence of the genes of interests were sequenced and only clones carrying homozygous frame-shifts leading to an early stop codon were kept. In order to generate a combined ZO1/ZO2 knock down KD line, we first created a ZO1 knock down and then we targeted ZO2. The ZO1 KD clone was mutant for two alleles, both alleles have a 1 bp insertion in the guide region (ACACACAGTGACGCTTC-1 bp insertion-ACAGGG) leading to an early stop of translation. The ZO2 KD has 5 bp deletion at the end of the guide region (GTACACTGTGACCCTACA-5 bp deletion-GG) leading to an early stop. Immunostaining and western-blot analysis showed that for ZO1 and ZO2 there was a residual expression of 2 - 3% of the wild-type levels present. We speculate this could be caused by rare ribosome mistakes at the indel mutations causing a back-in-frame shift (Makino et al., 2016).

To generate a N-terminal mNeonGreen knock-in (KI) the initial exon of ZO1 and ZO2 was targeted in the same region of the exon using the same guides as for KD. The gRNA and the trans-encoded RNA was annealed with Cas9 to form the riboprotein. The riboprotein and the repair plasmid were co-transfected *via* electroporation and overlap with the exon sides. Electroporation of each complex was performed in 300.000 cells (Invitrogen NEON electroporation machine and kit, 2 pulses, 20 ns, 1200 V). Cells were recovered 48 h after the electroporation. NeonGreen positive cells were selected by FACS 1 week after transfection. Correct insertion of mNeonGreen was verified by whole sequencing the ZO1 and ZO2 loci.

##### 3D cell culture and trans-epithelial permeability assay

To grow 3D MDCK-II cysts, the surface of glass coverslides (thickness 0.17 mm) was coated with a solution of laminin 0.5 mg/ml for 1 h at 37 °C 5% CO2. Cells were re-suspended from a 2D monolayer by action of 0.25% Trypsin EDTA for 10 minutes at 37°C 5% CO2 and a single cell suspension of 16.000 single MDCK-II cells per cm2 was seeded on the coated surface in complete media MEM, 5% v/v FBS, 1% v/v NeAA, 1% v/v Sodium pyruvate, 1% v/v Glutamax, 1% v/v Penstrep complemented with 15% Matrigel (Corning MG matrix, ref. 356231). Cells were cultured for 4 to 5 days in 37°C 5% CO2 until reaching 30 to 40 μm in diameter. To measure permeability of dextran (Alexa647, 10k) and a lipid tracer (DPPE-TMR) cyst medium was supplemented with 1μM DPPE-TMR, which was complexed with fat-free BSA in 1:1 molar ratio, for 8min at 37°C. After washing, the medium was supplemented with 10μM dextran. After 15min incubation cyst were imaged on a confocal microscope to evaluate distribution of the lipid tracer and dextran. Dextran permeability into the lumen was quantified by measuring the intensity in the lumen and dividing this to the average intensity outside the cysts. In the way, permeability of the lipid tracer was measured by normalizing the intensity at the apical membrane to intensity at the basal membrane to cyst.

